# Metabolic brain networks switch between a sparsely connected baseline and highly integrated states to support cognition

**DOI:** 10.64898/2025.12.17.695042

**Authors:** Hamish A. Deery, Emma Liang, Chris Moran, Gary F. Egan, Sharna D. Jamadar

## Abstract

The human brain achieves cognitive flexibility by rapidly switching between large-scale functional network states. While state switching is assumed to be energetically demanding, the direct metabolic foundation underpinning state switching has not been characterised. Here, we combine functional FDG-PET (fPET) imaging and sliding-window analyses to characterise metabolic network state switching in 85 adults (20-89 years). Four recurring metabolic network states were identified: a highly prevalent, sparsely connected baseline alongside three transient, globally integrated network states of cognitive control and attention. Cognitive performance relied on the capacity to mobilise the integrated associative system states. Flexible network switching was enabled by the moment-to-moment dynamic range of the underlying regional glucose signals. Neurotransmitter analyses revealed a low-dimensional neurochemical hierarchy that mapped to the dynamic network states, anchored by a dominant axis of endocannabinoid, metabolic, serotonergic and GABAergic systems. The dynamic metabolic network architecture was also attenuated in older adults, who exhibited a loss of network flexibility and were anchored to the sparsely connected baseline state. These findings reveal that functional network state switching is an emergent property of the brain’s metabolic architecture. This understanding may provide critical new insights into the metabolic basis of brain ageing, neurodegeneration and psychiatric conditions.

## Introduction

The human brain is a highly dynamic system that achieves adaptive behaviour by rapidly reconfiguring its large-scale functional network architecture between states of integration and segregation [1–6]. These dynamic transitions are believed to be metabolically expensive [7–9], requiring significant energy to fuel local network communities, engage high-cost central hubs and maintain long-range integrative pathways during state switching [7, 10–13]. An optimal balance is also required between network state switching and stability to enable the flexibility needed to adapt to new demands together with the consistency required to sustain attention [2, 4]. This balance is altered in ageing [14] and disrupted in neuropsychological conditions [15], leading to cognitive decline and neurodegenerative illness.

Functional networks form when spatially distributed populations of neurons exhibit coherent activity over time scales ranging from seconds to hours [5, 16, 17]. Consequently, network states and large-scale information processing emerge from the transient coalescence of neuronal firing patterns into rich, complex structures within an optimal dynamic range [7, 18–28]. It is well established that the brain’s electrophysiological and hemodynamic systems exhibit these network state characteristics to mediate cognition [29–31]. Recent discoveries indicate that the brain’s metabolic system exhibits a functional network architecture that is similar to, yet distinct from, the organisation of electrophysiological and hemodynamic networks [11, 32–36]. Characterised by frontoparietal predominance [11, 32] and macro-scale gradient characteristics [37, 38], this metabolic network architecture is predictive of higher-order cognition [34, 39], is grounded in genetic metabolic signalling [37] and is variable across the normative adult lifespan [11, 33]. This metabolic network architecture is also enabled by a cerebral *glucodynamic* system that is maintained in an optimal dynamic range [39].

It is unknown whether metabolic dynamics coalesce into transiently stable network states. Resolving this question is important to our understanding of the role of coherent cerebral metabolism in the emergence of cognition and complex neuropsychological phenotypes. Furthermore, addressing this question may provide crucial foundational evidence establishing cerebral metabolism as a complex and dynamical system that exhibits *metastability* [4], a leading computational model for explaining how cognition *emerges* from neural dynamics [40].

The gap in our understanding of metabolic network state switching has stemmed largely from the use of steady-state statistical models to measure brain metabolism [41] and technical limitations in measuring the moment-to-moment variability in cerebral glucose metabolism in the human brain *in vivo* [7]. The primary method for studying glucose metabolism in the brain *in vivo* is positron emission tomography with 18F-fluorodeoxyglucose (FDG-PET) [42]. Recent breakthroughs in time-resolved functional FDG-PET (fPET) have overcome the historical technical constraints and now make it possible to track rapid fluctuations in cerebral glucose metabolism within individual people [32, 36, 43–46], a phenomenon we refer to as "glucodynamics" [7, 39]. Because glucodynamics directly index postsynaptic metabolic flux, they provide a direct measure of the energetic substrate supporting neural signalling [7, 32, 47]. While correlations among regional glucodynamic signals define a large-scale metabolic network of the brain [11, 32, 34, 35, 39, 44], all work in metabolic connectivity to date has modelled connectivity across the full scan duration, effectively treating the network properties as static [20]. To determine whether metabolic network organisation dynamically reconfigures, it is necessary to apply time-variant methodologies to study metabolic connectivity.

To our knowledge, this study is the first to investigate time-variant brain network dynamics within the metabolic domain by extending "chronnectomic" [17] approaches to FDG-PET data. To help guide our methods and hypotheses, we propose a conceptual framework that draws on resource constraint principles [9, 48] and theories of coordinated dynamics [3, 49], economical brain organisation [50–53] and self-organised neural dynamics [54, 55]. Within this framework, the variability and complexity of underlying regional signal dynamics serve as the fundamental drivers of large-scale network state switching. Because the brain possesses limited internal glucose storage, it relies on a scalable, on-demand glucose supply via the vasculature ([39] also see [7, 48, 56]). We reason that the coupling between this dynamic but limited glucose supply and neural activity acts as a self-organising constraint on brain dynamics, regulating when and where neural activity can occur, its coordination in neural circuits and the balance between network integration and segregation [2, 3, 7, 14, 48]. In other words, the variable and complex structures embedded within these underlying metabolic signals act as a self-organising constraint, regulating the magnitude, speed and diversity of network state switching.

Our goal was to characterise metabolic network state switching and elucidate its relationship with cognitive performance and ageing. We hypothesised that better cognitive performance is associated with more dynamic metabolic network state switching. We also hypothesised that variable and complex regional glucodynamics enable a broad repertoire of centrally connected network states. We further hypothesised that metabolic network state switching declines in ageing.

## Results

We investigated metabolic network state switching using fPET data from 85 healthy adults in the Monash *MetConn* dataset [11, 33], including 40 younger (aged 20 to 42 years) and 45 older (66 to 89 years) people (see Supplement 2.1 for demographics). Participants completed a simultaneous 90-minute resting-state functional FDG-fPET-fMRI scan, and underwent a battery of cognitive assessments measuring memory, processing speed, response inhibition and task switching.

We used sliding window analyses to measure the time-variance and state switching characteristics of the metabolic network. Our primary analysis used windows of four, 16-second fPET frames (1:04-minutes). There was a 50% overlap with the preceding and following window. Sensitivity analyses using other window lengths are also reported in Supplement 2.7 and are described briefly below.

To characterise the recurring metabolic network states over time, 100 x 100 region [57] time-varying connectivity matrices were vectorised and clustered across the entire cohort using the K-means algorithm. Four distinct metabolic states (k = 4) were identified and their temporal dynamics measured using: 1) the total *number of transitions* to capture the global frequency in network switching; 2) *fractional occupancy rate* to measure the percentage of total windows a participant spent within a given network state; and 3) *mean dwell time* reflecting the average number of consecutive windows a participant remained within a given state configuration during a single uninterrupted visit.

To evaluate the topological architecture of the four metabolic network states, graph-theoretical metrics were calculated. Clustering coefficient was used to evaluate local network segregation and nodal strength centrality was calculated for each region to indicate its role as hub of central communication in the network [10]. To determine the metabolic cost associated with these topological network characteristics, we calculated a Glucose Cost Index (GCI) for the four network states, computed as the ratio of a region’s combined topological score - the geometric mean of its normalised clustering and centrality to its regional glucose consumption (CMR_GLC_). This metric quantifies ’metabolic cost’ by capturing the topological return per unit of baseline glucose [11, 52], revealing whether certain states are energetically optimised for information processing.

We begin by characterising the four metabolic network states before evaluating how their switching dynamics - specifically transition frequency, fractional occupancy, and dwell time - associate with cognitive performance. We then quantify the energetic profiles of these configurations using the GCI. To determine the signal dynamics associated these network shifts, we examine the relationship between state switching and the variability (SD) and complexity (entropy) of the underlying fPET timeseries. We also examine the neurochemical profiles of each state by spatially aligning regional centrality with neurotransmitter receptor maps. Finally we test for age-group differences across the network switching metrics.

### Characterising Metabolic Network State Switching

Cluster validity criteria evaluated across k = 2-8 did not reveal a sharp or unequivocal inflection point based on Silhouette width and within-cluster sum of squared errors (WCSS) elbow curves (see Supplement 2.2). Furthermore, at k = 4, clustering yielded a low mean Silhouette score that was not significantly higher than block-bootstrapped surrogates. These results align with established dynamic network connectivity literature in both fMRI [58, 59] and electrophysiological studies [60]. Because time-resolved functional network connectivity operates in a high-dimensional feature space (thousands of network edges), individual windowed matrices tend to occupy a smooth, continuous manifold rather than forming isolated cluster islands [20, 61].

Importantly, block-bootstrap surrogate testing (n = 100) confirmed that the k = 4 model captured a non-random dynamic structure, yielding a significantly lower WCSS in the real data compared to phase-shuffled surrogates (p = 0.009; Cohen’s d = -71.7). This significantly lower WCSS confirms the model’s validity in summarising primary metabolic network configurations within a continuous connectivity landscape. Selecting k = 4 for our primary analysis is also consistent with empirical precedents in neuroimaging studies, where k = 4–7 states capture macroscale dynamic reconfigurations without over-fractionating the time series into noisy sub-states [17, 58, 62, 63].

The temporal and network characterisation of the four states revealed salient differences in their occupancy and topological architecture (Figure 1). State 1 captured 17.7% of windows and demonstrated a network structure with moderate clustering (0.08) and centrality (8.19). State 2 served as the second most frequent network state, accounting for 18.0% of the total windows. Like State 1, State 2 exhibited intermediate clustering (0.06) and centrality (6.40). State 3 represented an infrequently visited but highly integrated state, appearing in 6.5% of windows. Despite its low occupancy, State 3 exhibited very strong global integration and local community structures, with centrality (20.34) and clustering (0.20) reaching their highest levels. Finally, State 4 was by far the most persistent configuration, capturing 57.8% of windows. This highly prevalent state was characterised by low centrality (1.60) and clustering (0.01), indicating a sparsely connected metabolic state. Moreover, the topology of State 4 was strongly segregated between the left and right hemispheres. Together, these features of State 4 suggest that it is a low-coherence, high occurrence baseline state. Similar results have been reported in resting state functional connectivity dynamics, where connectivity patterns are evident in specific states for long periods [58].

**Fig 1.**
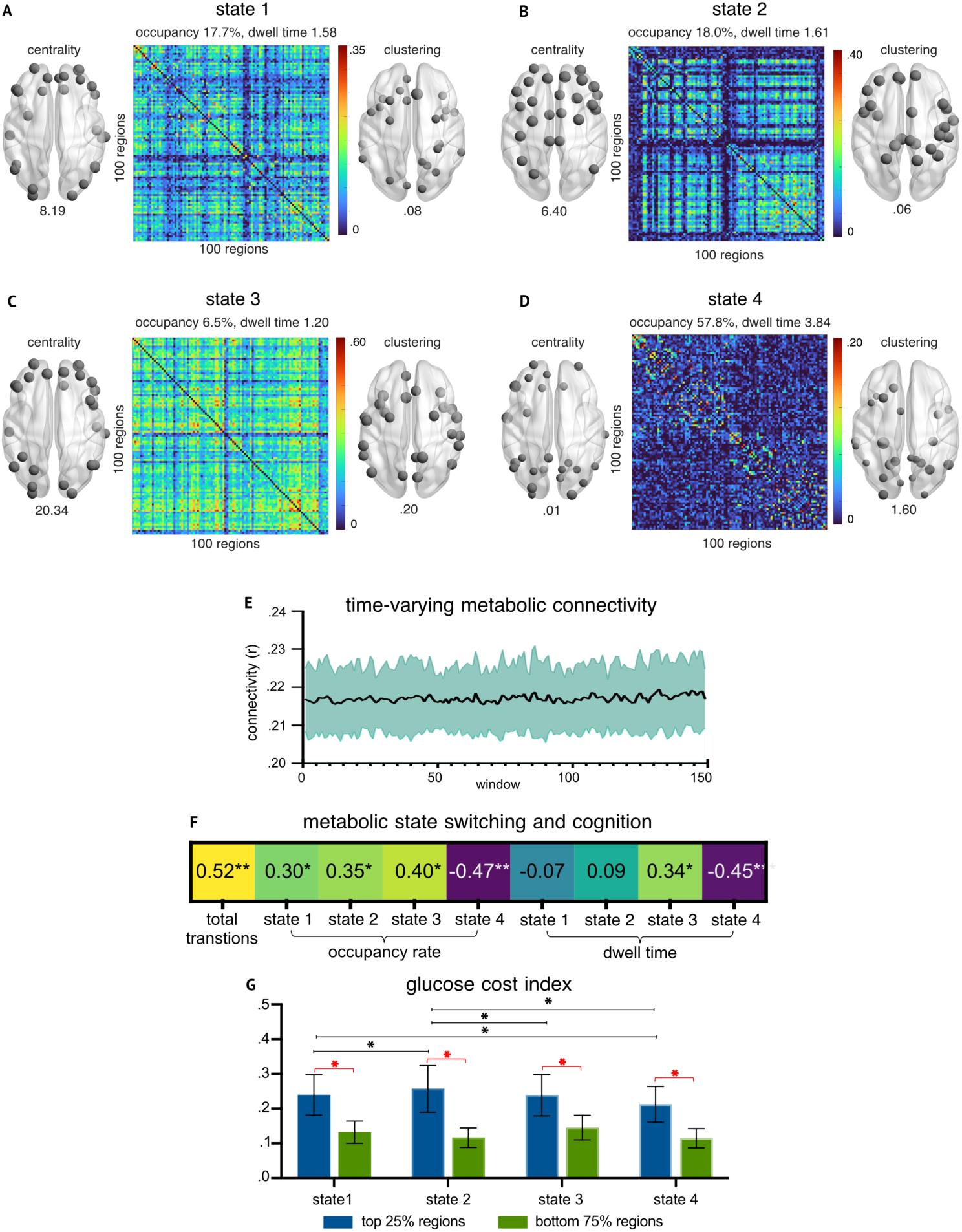
Metabolic network state switching. Four metabolic network states (A-D) were identified from K-means clustering of the metabolic connectivity timeseries (E; windows are four 16 second frames). For A-D, the connectome is shown for each of the four network states, together with the 25% of regions with the strongest centrality and clustering projected to the brain surface to illustrate regions most prominent in each network state. The numbers under the brain images represent the mean graph metric values to illustrate the relative strength across states (note: within each state, node size was auto-scaled to support visualisation). Association (Pearson correlations) between metabolic network state switching measures and cognitive performance (F). Cognitive performance is a single principal component including Hopkins Verbal Learning Test, category switch, digit symbol substitution and stop signal task performance. Stop signal and category switch reaction times were multiplied by -1 so that greater scores reflect better performance (**p-FDR <0.01, *p-FDR<0.05). Glucose cost index (GCI) across the four states (G), calculated as the ratio of combined topological score (geometric mean of normalised clustering coefficient and centrality) to regional CMR_GLC_. Within each state, the top 25% of regions based on combined topological score showed significantly higher GCI compared to the remaining 75% of regions (red asterisks). Significant differences in GCI were also observed across states (p < 0.001), with Bonferroni-corrected post-hoc comparisons showing State 2 had higher GCI, than all other (black asterisk p < 0.001). State 1 and State 3 showed comparable GCI, while State 4 demonstrated significantly lower GCI than all other states.

To characterise the topology of the network states, the top 25% of regional graph metrics were assessed and mapped to the brain surface (Figure 1A-D; also see Supplement 2.3), revealing distinct integration profiles and hub configurations. State 1 represented an associative system and visual integration profile, with prominent clustering and central hubs in lateral, posterior and dorsal prefrontal areas in the salience ventral attention, default and control networks, alongside visual and temporal regions. State 2 emerged as a coordinated associative system and somatomotor configuration, driven by regions of control and default mode networks; here, local clustering was anchored in lateral prefrontal control regions, while high-throughput centrality was dominated by prefrontal and temporal default nodes and somatomotor regions. The presence of regions from the same associative networks across States 1 and 2 is consistent with fMRI studies reporting that connectivity patterns of frontoparietal regions can change within and across states as the brain dynamically reconfigures it networks [13, 53, 64].

State 3 was characterised by a high-throughput core of global integration across associative networks, with seven of the top eight hub regions in the prefrontal cortex - spanning the control, default, salience and dorsal attention networks. This high-throughput core was supported by local clustering of regions anchored within attention networks (including the salience network’s insular nodes) alongside primary visual and somatomotor regions. Conversely, State 4 functioned as a low-coherence, hemisphere segregated baseline, characterised by relatively low graph metric values across all regions.

Together, these four states reflect a dynamic metabolic network topology. The metabolic network switches between a sparsely connected baseline and configurations of highly integrated regions supporting global communication. Crucially, the profiles of these states highlight a trade-off between temporal persistence and topological diversity. The clear dominance of the sparsely connected baseline (State 4) paired with the infrequent, punctuated appearances of the moderate (States 1 and 2) and highly (State 3) integrated states suggests that time-variant metabolic connectivity balances a highly stable yet diverse repertoire of network configurations. While the network spends the majority of time anchored in the high-occupancy baseline, it retains the capacity to dynamically traverse a range of highly-connected topological boundaries. These configurations indicate that while metabolic dynamics exhibit prolonged periods of temporal stability, the underlying architecture successfully accesses a rich and diverse range of functional network states.

### Metabolic Network State Switching and Cognition

To test our hypothesis that better cognitive performance is associated with more dynamic metabolic network state switching, we first reduced the dimensionality of the cognitive data using Principal Components Analysis. One significant PC was identified (Eigenvalue > 1), explaining 54.6% of the variance. The loadings for the cognitive measures were: HVLT (0.53), category switch (0.54), digit symbol substitution (0.55) and stops signal (0.36).

We next evaluated the correlations between the cognitive PC and the network state switching measures, revealing a distinct pattern of associations (Figure 1F). A greater number of total transitions between network states was associated with better cognitive performance (r = 0.52, p-FDR< 0.001). When evaluating individual network configurations, opposing directional profiles emerged based on state topology. For the integrated associative and attention network states, higher occupancy rates were associated with better cognitive performance for State 1 (r = 0.30, p-FDR = 0.007), State 2 (r = 0.35, p-FDR = 0.003) and State 3 (r = 0.40, p-FDR = 0.001). For dwell time, only the highest integrated State 3 demonstrated a significant positive relationship with cognition, such that longer dwell times were associated with better cognitive performance (r = 0.34, p-FDR = 0.003). Conversely, greater affiliation with the sparsely connected baseline (State 4) was detrimental to cognitive performance; a higher occupancy in State 4 (r = -0.47, p-FDR < 0.001) and extended dwell times (r = -0.45, p-FDR < 0.001) were both associated with worse cognitive performance.

To isolate the specific temporal properties most driving cognitive performance, we ran linear multiple regression models predicting the cognitive PC from the occupancy rates and dwell times of the metabolic states. For occupancy rates - which are compositional and sum to 100% per subject - State 4 was omitted as the reference category to prevent perfect multicollinearity. For dwell times - which are not compositional - all four states were included in the model.

The state occupancy regression model explained 26.9% of the variance in cognitive performance (F(3, 73) = 8.9, p < 0.001). Switching out of the State 4 baseline into the highly integrated State 3 significantly predicted cognitive performance (ß = 0.37, p = 0.016). A moderate statistical trend was also observed for switching into State 2 (ß = 0.22, p = 0.074). The dwell-time regression model was also significant (F(4, 73) = 6.3, p < 0.001), explaining 25.8% of cognitive variance. The mean dwell time of the baseline State 4 emerged as the sole predictor of cognitive performance, with higher dwell time associated with worse cognitive performance (ß = -0.46, p = 0.002). These regression models indicate that cognitive performance relies on the brain’s capacity to reconfigure its metabolic network architecture into integrated states, particularly State 3, before switching back to the baseline state.

Overall, these results demonstrate that metabolic network state switching during a non-task state is associated with cognitive performance. A higher capacity for complex information processing is associated with the capacity to switch between a sparsely connected baseline configuration and a flexible metabolic network, characterised by more frequent transitions between states and sustained activation of highly integrated associative and attentional networks.

### Glucose Cost Index for Metabolic Network States

To determine the metabolic cost associated with these network state transitions, we compared their Glucose Cost Index (GCI) scores. Higher GCI indicates a greater topological return (stronger regional clustering and/or centrality) for a given baseline glucose budget (CMR_GLC_). Within each network state, the top 25% of regions measured by clustering and centrality consistently showed higher GCI values compared to the bottom 75% of regions (all p < 0.001, Cohen’s d ranging from 3.55 to 4.03), indicating that regions with greater topological importance are less metabolically costly (Figure 1G). In other words, the most strongly connected regions in each network state achieve that connectivity at a lower per-unit glucose consumption compared to other regions.

A repeated measures ANOVA demonstrated significant differences in GCI values of the top 25% regions measured by clustering and centrality across the four metabolic network states, (F(3,237) = 261.55, p < 0.0001). Bonferroni-corrected post-hoc comparisons revealed that State 2 exhibited the highest GCI (0.257), significantly higher than State 4 (p < 0.001), State 1 (p < 0.001) and State 3 (p < 0.001). Notably, State 1 and State 3 showed comparable GCI (p = 0.173), despite their different topological profiles - State 1 being moderately integrated (centrality = 8.19, clustering = 0.08) and State 3 being highly integrated (centrality = 20.34, clustering = 0.20). State 4 consistently demonstrated significantly lower GCI than all other states, indicating that the most frequently attained baseline state has a high metabolic cost, yielding low local and global connectivity strength per unit of glucose consumption.

Together, these results demonstrate the metabolic network states vary in their efficiency profiles. Specifically, they suggest that the high occupancy baseline state has a high metabolic overhead, whereas the integrated states efficiently maximise the network and computational return on energetic investment.

### Glucodynamics and Metabolic Network State Switching

We next evaluated whether the underlying regional dynamics of the glucose signal govern the flexibility of metabolic network state switching. We reasoned that these transitions are energetically demanding and constrained by underlying regional glucodynamics, particularly when involving high-cost central hubs [7, 10, 12, 65]. Hence, we tested the hypothesis that glucodynamics act as the primary enabler of state switching between the highly integrated network states.

To quantify the glucodynamic properties we calculated the regional standard deviation and entropy of the fPET time series for each participant. While these neuroimaging signals both describe aspects of brain variability, they capture different properties (see Supplement 2.4 for illustration) and have each been associated with cognitive performance and ageing in fMRI and EEG studies [21, 24, 28, 30, 66, 67]. We calculated a glucodynamic index (average of z-scored regional sample entropy (fPET_EN_) and standard deviation (fPET_SD_) of the fPET timeseries across the 100 regions). Spatial characterisation revealed that fPET_SD_ was greatest in superior fronto-central regions and lowest in temporal cortices, whereas fPET_EN_ peaked in temporo-limbic regions and was lowest in parietal-central areas (Figure 2A-B).

**Fig 2.**
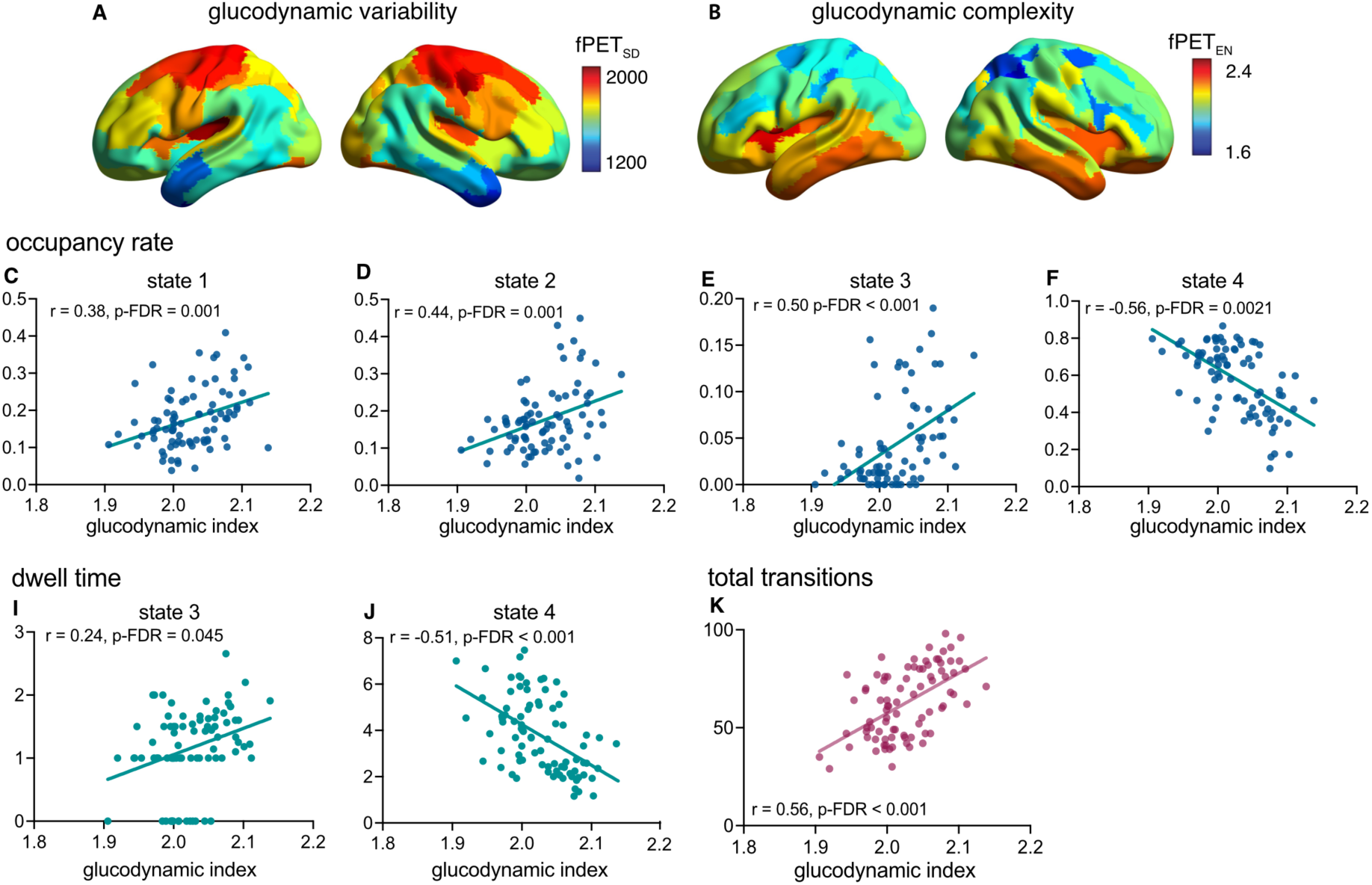
Metabolic network state switching and glucodynamics. Mean regional variability (fPET_SD_) (A) and complexity (fPET_EN_) (B) of the regional glucodynamic signals. Partial correlations between metabolic network state switching measures and glucodynamic index, controlling head motion (C-K). Glucodynamic index is the mean of the z-scored regional variability (SD) and complexity (entropy) for each participant. A higher glucodynamic index was associated with higher occupancy in states 1-3 and higher dwell time in state 3 (associations with dwell times in states 1 and 2 were not significant and are not shown, r = 0.09, p-FDR = 0.457 and r = 0.21, p-FDR = 0.079). In contrast, a higher glucodynamic index was associated with lower occupancy and dwell times in state 4 (F and J).

We next used partial correlation analyses to assess the association between the glucodynamic index and the metabolic state switching measures, controlling head motion. We observed robust relationships (Figure 2C-K). A higher glucodynamic index was associated with a greater number of total transitions between network states (r = 0.56, FDR < 0.001). Greater glucodynamics were also associated with specific temporal properties across the metabolic network states. For the integrated states (States 1-3), a greater glucodynamic index was associated with higher occupancy rates in State 1 (r = 0.38, FDR = 0.001), State 2 (r = 0.44, FDR = 0.001), and State 3 (r = 0.50, FDR < 0.001). For dwell time, greater glucodynamics were associated with longer periods in State 3 (r = 0.24, FDR = 0.007); but not States 1 and 2 (both p > 0.079). Crucially, this relationship inverted when evaluating the sparsely connected baseline state. A higher glucodynamic index was associated with lower State 4 occupancy (r = -0.56, FDR < 0.001) and dwell times (r = -0.51, FDR < 0.001). Together, these results indicate that more variable and complex glucose metabolism enables the flexibility of the metabolic network, appearing to drive the brain away from a prolonged, low-coherence baseline configuration toward a richer repertoire of globally integrated associative system states.

To quantify the specific underlying metabolic drivers of network state switching, we ran multiple linear regression models predicting the glucodynamic index from the state switching measures. The model utilising state occupancy rates (with the State 4 baseline again omitted as the reference category and head motion included) explained 31.2% of the variance in glucodynamics (F(4, 74) = 8.7, p < .001). Both higher State 2 occupancy (ß = 0.31, p = 0.009) and State 3 occupancy (ß = 0.27, p = 0.066) were positive predictors of increased glucodynamics. The model for dwell time explained 29.6% of the variance (F(5, 74) = 6.2, p < .001). Dwell time of the baseline State 4 was the only significant predictor of glucodynamics, with higher glucodynamics associated with lower dwell time (ß = -0.58, p < 0.001). Collectively, these results suggest that a higher glucodynamic range enables the metabolic network to flexibly switch out of the sparsely connected State 4 baseline into highly integrated network states that support adaptive cognition.

Because both the glucodynamic index and the network state switching measures are derived from the same fPET data, there is an inherent potential for mathematical coupling. However, the glucodynamic index measures are univariate (evaluating localised, regional signal fluctuations), whereas the state switching metrics are bivariate (evaluating inter-regional coherence across time). Moreover, our results demonstrate that these measures are not simply biased toward uniform positive correlations. For example, a higher glucodynamic index was robustly associated with a reduction in baseline state occupancy and dwell time (r = -0.56 and r = -0.51, respectively). If the relationship were purely a mathematical by-product of shared data variance, we would expect a uniform directionality of effect across all network configurations rather than the state-specific, directionally opposing dynamics observed here.

### Metabolic Network States and Neurotransmitter Associations

To gain a more mechanistic understanding of the four metabolic network configurations, we next characterised their neurochemical profiles leveraging the *neuromaps* toolbox [68]. Regional centrality of the four metabolic states was spatially co-registered against an extensive *in vivo* repository of neurotransmitter receptor and transporter maps acquired from healthy cohorts using high-resolution positron emission tomography (PET) and single-photon emission computed tomography (SPECT). Because our network metrics were calculated using a purely cortical parcellation (Schaefer 100), all volumetric molecular repository maps were projected to the surface mesh space (fsaverage). To prevent artifactual signal distortion from subcortical structures or regional dropouts, a shared logical mask was applied dynamically to isolate and evaluate only the overlapping, valid cortical vertices across both the connectivity and molecular datasets.

Spatial correlation coefficients (*r*) were computed between each network state and the 31 cortical molecular maps (see Supplement 2.5 for full list), with statistical significance assessed using permutation testing (n = 1000 permutations). Following FDR correction, 120 of 124 metabolic state-receptor pairs were significant (p-FDR < 0.05; Supplement Table S6). These analyses identified spatial covariation between our empirically-measured metabolic connectivity states and independently-derived spatial maps of chemoarchitecture.

While individual molecular map-metabolic network state correlations provide narrow molecular insights, individual neurotransmitter systems exhibit strong spatial overlap and collinearity due to shared cellular distribution profiles and cortical hierarchies [69]. Hence, to extract the broad, latent neurochemical axes that coordinate metabolic network dynamics across configurations, an unsupervised pattern decomposition was implemented via Principal Component Analysis (PCA) across the state correlation matrix, and the loading profiles of both the states and individual receptors were evaluated to interpret their functional architecture. We first provide a qualitative and quantitative description of the principal components, followed by an interpretation of the receptor-PC loadings.

### Metabolic Network State-Neurochemical Spatial Alignment

The PCA decomposition revealed an architecture where the first three principal components accounted for 99.75% of the total spatial variance across the receptor-metabolic network state dataset. PC1 was the dominant latent axis, explaining 86.25% of the total variance, while PC2 captured 11.81%, and PC3 accounted for 1.69%.

Evaluation of the network state loadings across these axes revealed highly distinct mapping. PC1 demonstrated uniformly high, positive loadings across all four dynamic metabolic network states: State 3 (0.985), State 4 (0.963), State 1 (0.924), and State 2 (0.836). This indicates that PC1 represents a dominant, universal neurochemical organisation shared globally across the entire metabolic state repertoire, spanning from the sparsely connected baseline (State 4) to the highly integrated prefrontal core (State 3). As such, PC1 can be interpreted as a shared neurochemical scaffold upon which all metabolic network states are superimposed, consistent with the known cortical gradient organisation of the human brain [70]. In contrast, PC2 segregated the associative-somatomotor configuration of State 2 at the positive pole (+0.546) from the associative-visual network of State 1 at the negative pole (- 0.370). Finally, PC3 differentiated the two topological extremes of the metabolic network state repertoire, separating the highly prevalent baseline State 4 (+0.198) from the low occurrence, globally integrated State 3 (-0.148).

### PC1 Loading Architecture: The Universal Cortical and Metabolic Infrastructure

The primary axis was strongly associated with all four metabolic states. To identify the molecular systems anchoring this primary axis, we evaluated the receptor spatial loadings on PC1 (see Figure 3 and supplementary Table S6). The positive pole of PC1 was heavily dominated by the endocannabinoid receptor OMAR (CB_1_, λ = 0.937) [71, 72], alongside global markers of baseline cortical function including cerebral blood flow (MEANCBF, λ = 0.787) [73] and histone deacetylase (MARTINOSTAT, λ = 0.783) [74]. Widespread associative cortical targets also loaded strongly on this axis, including serotonergic (5-HT_6_, λ = 0.779; 5-HT_2A_, λ = 0.769; 5-HT_1A_, λ = 0.696) [75], GABAergic (GABA_A_, λ = 0.701) [76, 77], muscarinic M_1_ (LSN3172176, λ = 0.750) [78], and synaptovesicular density (UCBJ, λ = 0.677] [79] markers. Conversely, the negative pole of PC1 was associated with markers of subcortical modulatory systems, including histamine H_3_ (GSK189254, λ = -0.759) [76], μ-opioid receptor (CARFENTANIL, λ = -0.649) [80], noradrenaline transporters (MRB, λ = -0.643) [76], and dopamine transporters (DAT, λ = -0.616) [81].

**Fig 3.**
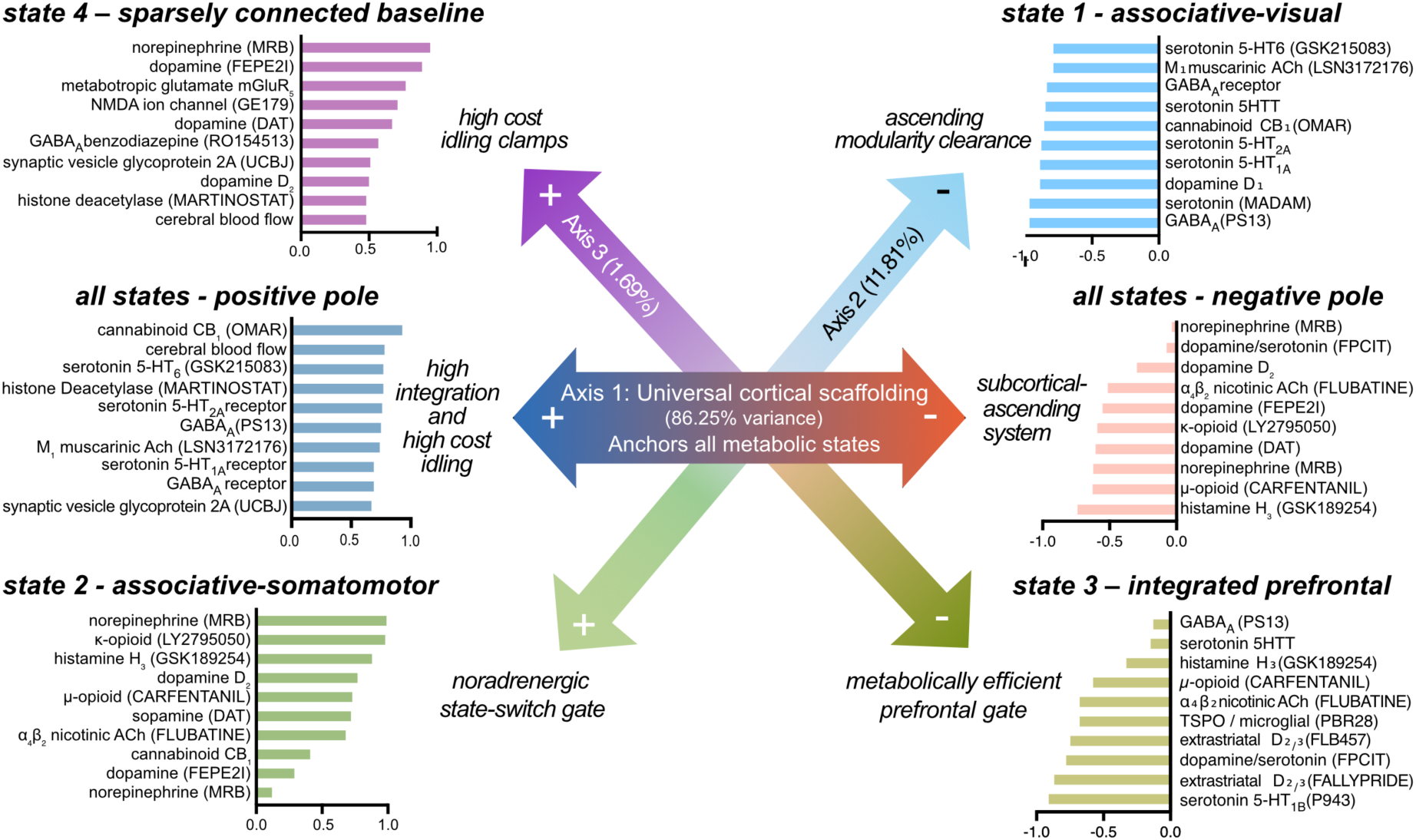
Low-dimensional neurochemical architecture governing dynamic metabolic brain states. A schematic representation of the three principal component (PC) axes capturing 99.75% of the spatial variance across the metabolic network states and 31 neurotransmitter maps. The 10 radioligands/transmitters most strongly loading on each pole of the axes are shown (see Supplement for full list). The horizontal axis (PC1, 86.25% variance) defines a universal cortical infrastructure shared across all four metabolic network configurations, contrasting a transmodal, high-throughput associative pole (+; anchored by endocannabinoid, serotonergic, and GABAergic systems) against a baseline modulatory pole (-; driven by subcortical-ascending modulatory footprints). The dynamic departures from this universal template are mediated by two orthogonal axes (diagonals) acting as specialised gating mechanisms. PC2 (11.81% variance) acts as a noradrenergic state-switching gate that segregates the active associative-somatomotor configuration of State 2 (+) from the associative-visual network of State 1 (-). PC3 (1.69% variance) functions as a fine-grained integration gate, separating the high-occupancy, high-cost baseline of State 4 (+) from the highly integrated, metabolically efficient prefrontal core of State 3 (-).

Together, these results suggest that all four metabolic network states are associated with a shared synaptic-metabolic-neuromodulatory architecture. This architecture is synaptically dense, metabolically active, regulatory and plastic, and underpinned by serotonin, acetylcholine, GABA and endocannabinoid chemoarchitectures. It is also anticorrelated with dopamine- and noradrenaline-driven arousal, reward and action systems. This pattern is consistent with a network architecture that supports an integrative and stable function rather than reactive, alerting, and action-oriented function. The results suggest that transitioning across metabolic network states does not require switching this macro-scale neurochemical framework; rather, all states operate atop this shared baseline scaffold. By excluding the subcortical structures through our cortical mask, the clear dominance of widespread, transmodal receptors over ascending modulatory footprints on PC1 suggests that the primary axis captures the intrinsic metabolic and neurochemical architecture of the human cortex.

### PC2 Loading Architecture: The Noradrenergic State-Switching Gate

PC2 characterised the spatial divergence between State 1 and State 2. The positive pole of PC2 - which maps to the associative-somatomotor configuration of metabolic network State 2 - was almost perfectly aligned with the noradrenaline transporter MRB (λ = 0.999) [76] and the kappa-opioid receptor (LY2795050, λ = 0.990) [82], alongside significant contributions from histamine systems (GSK189254, λ = 0.890) [76], extrastriatal dopamine (D_2_, λ = 0.780; DAT, λ = 0.729) [81, 83], and μ-opioid targets (CARFENTANIL, λ = 0.740) [80]. Conversely, the negative pole of PC2 aligned with the associative-visual metabolic network State 1 and was dominated by serotonin transporters (MADAM, λ = -0.981) [84], cortical serotonin receptors (5-HT_1A_, λ = -0.901;5-HT_2A_, λ = -0.888; 5-HT_6_, λ = -0.804) [75], dopamine D_1_ profiles (λ = -0.904) [85], and GABAergic markers (GABA_A_, λ = -0.848) [76, 77, 86].

Together, these results suggest that metabolic network States 1 and 2 occupy an axis defined by a catecholamine-opioid-histamine enriched network involved in behavioural mobilisation, motivational salience, and arousal regulation. While axis 1 represents a balanced, integrative, and stable pattern of function, axis 2 represents a pattern that may help the brain detect salient events and prepare a response, especially under conditions of reward, threat, pain, stress or uncertainty. This dissociation between the positive and negative poles of the axes suggests that while State 1 is aligned to an integrated, serotonergic and GABAergic associative system, exiting this configuration to enter State 2 requires a localised, chemically driven state-switch. The near-perfect loading of cortical noradrenergic projection footprints (MRB) on the positive pole is consistent with the idea that ascending locus coeruleus modulation provides the necessary cortical gain adjustment to dynamically shift away from associative hubs toward primary motor networks.

### PC3 Loading Architecture: The High-Cost Integration Regulatory Gate

Although explaining a smaller fraction of spatial variance (1.69%), PC3 captured a specific biological and topological contrast between the high-occupancy, sparsely connected metabolic baseline (State 4) and the rare, highly integrated, low cost network state (State 3). The positive, baseline pole of PC3 was defined by concentrated monoaminergic markers, specifically norepinephrine transporters (METHYLREBOXETINE, λ = 0.960) [87] and dopamine transporters (FEPE2I, λ = 0.895) [81], alongside metabotropic glutamate signalling (mGluR_5_, λ = 0.777) [88]. The negative pole, which tracks the global integration of State 3, was anchored by serotonin 5-HT_1B_ (P943, λ = -0.919) [89] and extrastriatal dopamine receptors D_2/3_ (FALLYPRIDE, λ = -0.880) [90].

These results indicate that States 3 and 4 occupy an axis defined by a catecholamine-glutamate-serotonergic-dopamine chemoarchitecture involved in alerting, action readiness and affective/motivational processes. Because State 3 reflects prefrontal connectivity across control, default, and salience networks, axis 3 appears to act as a homeostatic regulatory gate. The strong anchoring of norepinephrine and dopamine transporters on the positive pole implies that active monoaminergic clearance patterns over the cortex serve to maintain the stable, hemisphere-segregated baseline of State 4. Only when this modulatory clamp is transiently altered can the network access the rare, highly integrated transmodal core of State 3.

### Metabolic State and Neurotransmitter System Summary

Collectively, the results of the neurotransmitter analyses reveal that the brain’s dynamic metabolic network repertoire is aligned to a dominant neurochemical axis. The vast majority of spatial variance (86.25%) is anchored to a universal cortical scaffold of endocannabinoid, metabolic, and transmodal receptor profiles (PC1), which include key serotonergic, GABAergic, and synaptovesicular density markers [69]. Dynamic departures from this shared axis appear to be achieved via a specialised noradrenergic state-switching gate (PC2) that dissociates somatomotor metabolic networks (State 2) from visual metabolic states (State 1), and a fine-grained modulatory gate (PC3) that regulates the transitions between the baseline metabolic network state (State 4) and a globally integrated prefrontal state (State 3). By flexibly shifting metabolic connectivity along these distinct, low-dimensional molecular axes, the cortical network successfully balances temporal persistence with a rich, flexible repertoire of functional topologies.

### Age Differences in Metabolic Network State Switching

As a final aim, we evaluated age-group differences in the metabolic network state switching measures using *t*-tests (Figure 4). Older adults exhibited a marked reduction in network flexibility, demonstrating significantly fewer total transitions between brain states compared to younger adults (t = 7.4, p-FDR< 0.001; Figure 4I). Ageing was also associated with a notable shift in the temporal profile of network state switching. For the integrated, higher-order networks states, older adults exhibited significantly reduced occupancy rates in State 1 (*t* = 5.2, p-FDR< 0.001) and State 2 (t = 3.4, p-FDR= 0.002). A similar age group difference was seen for high integration State 3 (*t* = 5.7, p-FDR< 0.001), with a notable profile of the large majority of older adults failing to occupy this state even infrequently, if at all (dots clustering around zero in Figure 4G). Similarly, the duration of individual visits to this highly integrated State 3 showed an age-related decline, with older adults having significantly shorter dwell times when they did switch to this state (*t* = 3.5, p-FDR= 0.001). Conversely, the metabolic system became anchored in the sparsely connected baseline state (State 4) in older adults. Older adults spent significantly more time in this configuration, demonstrating a substantially elevated State 4 occupancy rate (*t* = -7.2, p-FDR< 0.001) paired with significantly prolonged dwell times (*t* = -6.4, p-FDR< 0.001).

**Fig 4:**
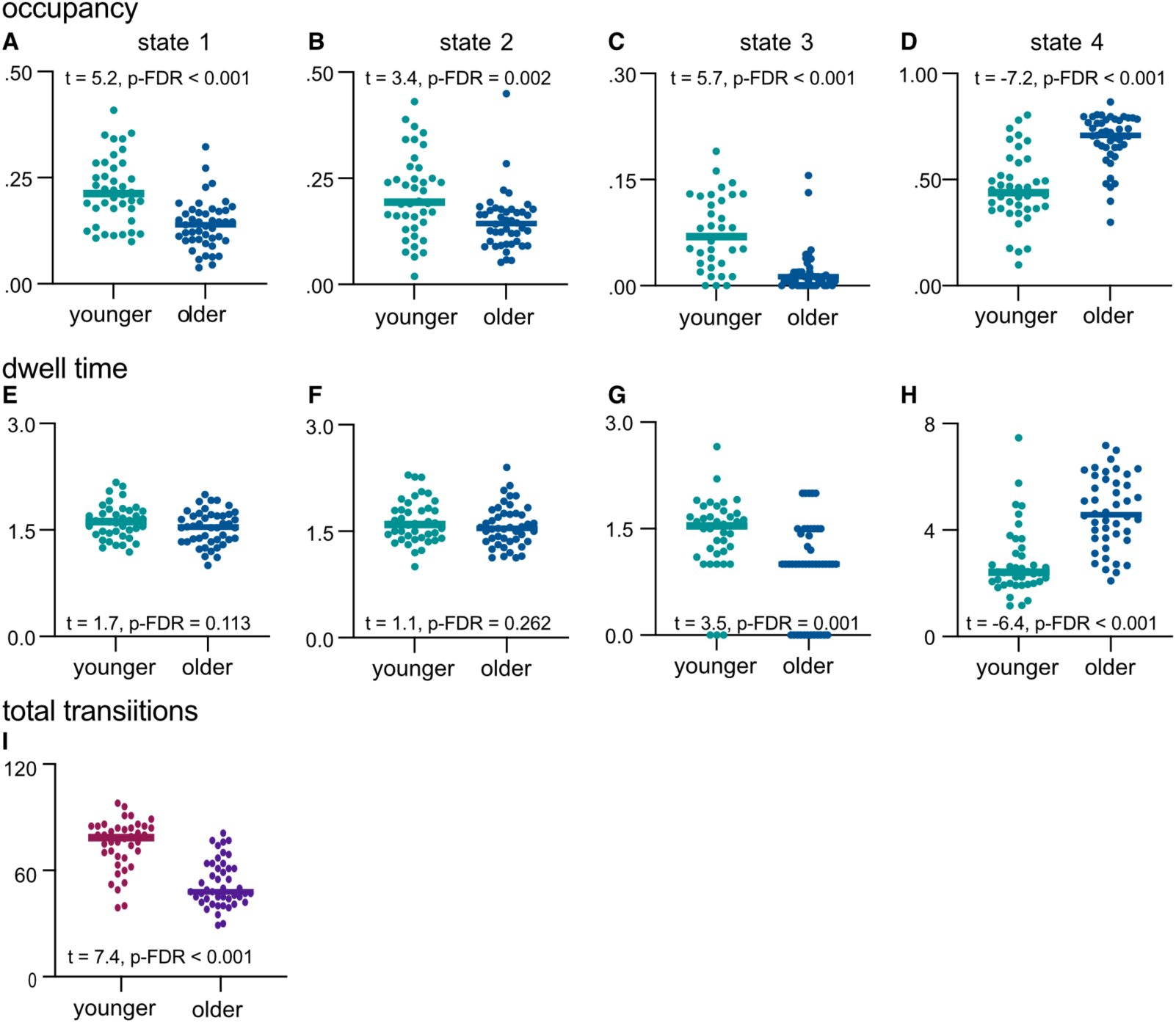
Age group differences in metabolic network state switching. Age group differences in metabolic network state switch measures (A-I). Older adults exhibited significantly reduced occupancy in States 1-3 (A-C), reduced dwell time in States 3 (G), and fewer total transitions (I). Conversely, older adults had higher occupancy and greater dwell time in State 4 (D and H). Solid line is median in A-I.

These findings are compatible with previous work demonstrating that healthy ageing is associated with overall reductions in global metabolic connectivity [11, 33]. Our results extend this framework to establish that ageing constrains metabolic network dynamics and state switching. Rather than flexibly traversing a diverse repertoire of globally integrated network states, the metabolic system in older people becomes anchored within a sparsely connected baseline configuration, reflecting a loss of global information processing capacity.

### Validation: Metabolic Network State Switching with Alternative Sliding Window Lengths

The 64 second duration used to compute the sliding window connectivity and network state switching measures is relatively long and was necessitated by the 16 second duration of the fPET frames. Nonetheless, to ensure that our observed metabolic network states were not artifacts of the four-frame sliding windows, we conducted a sensitivity analysis across alternative window lengths of six and eight frames (Supplement 2.7). Across these alternate temporal scales, the cluster validation metrics supported a four-state solution. Furthermore, the overall chronnectomic architecture remained robust: State 4 consistently emerged as the dominant, low-coherence baseline (∼54–59% occupancy) and displayed spatial profile stability across alterative window lengths (minimum r = 0.928). Conversely, State 3 consistently maintained its profile as a low occupancy, highly integrated network state. We also observed a proportional drop in mean transitions (62.9 to 32.3) and scaling of dwell times as window size increased, reflecting expected temporal effects of combining longer measurement periods (more windows) and confirming that the macroscale metabolic dynamics described in our primary analysis are methodologically and biologically sound.

## Discussion

Our results show for the first time that the brain’s dynamic energy landscape constrains its repertoire of network configurations. Previous metabolic connectivity work has relied on time-invariant, single measures of connectivity and network integration averaged over time [32, 34–36, 41]. By introducing a sliding-window approach, we demonstrate that the time-variant flexibility of the metabolic network is an important determinant of adaptive neural processing and cognitive performance. The brain dynamically shifts between distinct metabolic network states, transitioning along a smooth, continuous connectivity landscape and switching between a sparsely connected baseline (State 4) and configurations characterised by integrated frontoparietal control and attention hubs (States 1, 2 and 3). These rich metabolic network transitions are associated with variable and complex regional glucodynamics, better cognitive performance and attenuate in ageing. Metabolic networks map to a shared neurochemical architecture across states, alongside specialised noradrenergic state-switching and fine-grained neuromodulatory gates that enable the brain to attain globally integrated configurations. By switching metabolic connectivity along these distinct, low-dimensional molecular axes, the cortical network appears to successfully balances temporal persistence with a rich, flexible repertoire of functional topologies. Collectively, our results support the idea that brain metabolism is not merely a downstream metabolic correlate of neural activity but a key enabler of macroscale functional network dynamics and state switching [7, 39].

Metabolic network state switching is characterised by a trade-off between temporal persistence and topological variety. The majority of the metabolic time-course (58%) is spent in a sparsely connected, hemisphere-segregated state (State 4), representing a highly stable baseline configuration. Punctuating this baseline are transient excursions into highly connected associative and sensory states (States 1-3). The low-occurrence and most highly integrated State 3 (6.5% occupancy) functions as a high-throughput communication state, heavily driven by centralised prefrontal hubs across control, default, and attention systems, and neurochemically aligned with serotonergic feedback and dopaminergic modulation systems. Our findings reveal that this shifting landscape is functionally important; a higher frequency of total network transitions and a greater sustained occupancy within the integrated associative system states (States 1-3) are related to superior cognitive performance. Conversely, an inability to escape the sparsely connected baseline (State 4) is significantly detrimental to cognitive function.

Our Glucose Cost Index (GCI) analyses indicate that the baseline state is the most metabolically costly configuration, demanding high glucose consumption relative to its low topological return. This costly baseline state is consistent with work indicating that a high proportion of the brain’s metabolic “budget” is spent maintaining neural integrity and associated homeostatic processes, with transitions to task-evoked states requiring only incremental metabolic cost [52, 91–93]. State 3 achieves its high-throughput connectivity at a higher glucose cost than the moderately integrated State 2, which appears to occupy an energetic "sweet spot" that maximises functional output per unit of glucose. Together, these results demonstrates that cognition relies on a highly flexible metabolic network repertoire governed by the capacity to rapidly mobilise integrated and variably efficient associative system states away from metabolically expensive baseline activity.

The relationship between large-scale metabolic network flexibility and the underlying dynamics of the glucose signals offers a framework for understanding the neurological trajectory of healthy ageing. Consistent with fMRI and EEG studies of network state switching [26, 27, 94–96], older adults explore a less diverse range of metabolic network states than younger adults, demonstrating fewer network state transitions. Rather than flexibly traversing the network repertoire, older adults spend a high proportion of time anchored within the sparsely connected baseline (State 4). Concurrently, integrated associative configurations (States 1-3) show age-related declines in both occupancy rates and dwell times, underpinned by attenuated signal dynamics. Our results suggest that preserved metabolic variability and complexity within these key networks can support cognitive resilience in older people, maintaining the state-switching capacity required for effective cognitive processing.

The ability of the brain to dynamically shift between metabolic network states aligns with frameworks of resource constraint principles [9, 48] and theories of coordinated dynamics [3, 49], economical brain organisation [50–53] and self-organised neural dynamics [54, 55]. In these frameworks, efficient information processing requires the brain to perpetually balance a "push-pull" dynamic: segregating into specialised, local clusters for routine tasks and rapidly integrating global configurations to manage novel or complex environments [16, 50, 53]. Similarly, under a predictive processing framework, the brain works to minimise prediction error while navigating an uncertain environment. This balance requires the system to maintain *metastability* or the capacity for individual regions to simultaneously preserve local function while transiently coalescing into global networks [3, 4, 14, 97, 98]. Our GCI metrics provide direct empirical weight to these frameworks by revealing a spatial and temporal economy in the brain: across all configurations, the top 25% most topologically important regions consistently display significantly higher GCI values, demonstrating that high-throughput hubs achieve stronger connectivity at a lower per-unit glucose cost. The near-equivalence in GCI between the different topological profiles of State 1 and State 3 further underscores this dynamic flexibility, indicating that the brain can access a flexible repertoire of integrated configurations to maintain efficient processing across diverse cognitive demands before relaxing back into low-coherence baseline states.

Our results also align with theoretical perspectives that conceptualise the human connectome as traversing a non-linear energy landscape where configurations occupy distinct “attractor basins” [99, 100]. The dominant, sparsely connected baseline state (State 4; 58% occupancy) represents a highly stable equilibrium basin. Conversely, switching to infrequent, globally integrated configurations (State 3; 7% occupancy) represents an energetically demanding transition away from baseline, requiring the system to overcome activation barriers to achieve widespread integration [99]. Our findings suggest that a higher glucodynamic complexity expands the metabolic dynamic range, preventing the brain from becoming caught in the sparsely connected baseline attractor basin and enabling the updating of internal models to process complex environmental demands before switching back to the baseline state [55, 97, 101].

The bridge between macroscale metabolic network state properties and the underlying low-dimensional neurochemical hierarchy that scaffolds the human cortex offer three key advancements in our understanding of brain organisation. First, our findings reveal that the brain’s dynamic metabolic repertoire likely operates with molecular efficiency. The dominant association with a primary neurochemical axis (PC1, 86.25% variance) suggests that transitioning between distinct metabolic network states does not require a wholesale reconstruction or large-scale spatial rearrangement of the brain’s neurochemical profile. Instead, all dynamic configurations - spanning from a sparsely connected baseline to a globally integrated prefrontal core - operate atop a universal cortical neurochemical backbone. This primary infrastructure establishes a baseline "push-pull" relationship across the cortex, balancing highly localised, transmodal processing (endocannabinoid, serotonergic, and GABAergic receptor systems) against ascending modulatory clearance footprints (histamine, μ-opioid, and noradrenergic systems). The human cortex utilises this baseline template to preserve network stability while executing fluid, state-to-state dynamic transitions.

Second, this work extends traditional models of how neurotransmitter systems dictate large-scale network function. While classical neurochemistry often treats modulatory systems as independent diffuse systems [102], our low-dimensional decomposition suggests that these systems may act as coordinated, highly specific topological "gates" to manage the brain’s macroscale landscape. A specialised noradrenergic switch (axis 2) appears to enable the brain to exit a stable, integrated associative-visual regime to mobilise primary motor networks. The near-perfect alignment of cortical noradrenergic projection footprints on this axis provides evidence that ascending locus coeruleus inputs act as a centralised gain-control mechanism to rapidly shift the brain’s global metabolic topology. Furthermore, a fine-grained modulatory gate (axis 3) seems to regulate access to the low occurrence, metabolically integrated prefrontal core. The heavy concentration of monoaminergic transporters on this axis suggests that active cortical clearance of norepinephrine and dopamine behaves as a protective chemical "clamp," deliberately maintaining the brain in the baseline state. Although this baseline state demands a higher metabolic overhead relative to its modest topological connectivity, this costly baseline equilibrium may act as an essential computational safeguard, preventing excessive global synchronisation. Widespread network integration is only achieved when this modulatory clamp is transiently altered by distinct serotonergic (5-HT_1B_) and extrastriatal dopamine (D_2/3_) profiles.

While speculative, this multi-axis molecular hierarchy has implications for the attenuation of glucodynamics observed in ageing. Because the transition out of the sparsely connected baseline (State 4) into the integrated prefrontal core (State 3) or active motor networks (State 2) depends on the precise mobilisation of ascending noradrenergic (PC2) and monoaminergic clearance (PC3) mechanisms, the age-related reduction in state transitions likely reflects a structural or functional deterioration of these modulatory gating systems [103]. When local glucodynamics flatten in older people, the system loses the energetic capacity to engage these specialised chemical switches, ultimately anchoring the ageing brain within the low-coherence baseline configuration.

To build upon this framework, the next step in this research is to transition from static, *in vivo* molecular atlases to active pharmacological or task-based interventions combined with dynamic functional PET. Because the data used to construct these neurochemical axes reflect group-level receptor densities [69, 104], future studies should employ targeted pharmacological challenges (e.g., administering noradrenergic reuptake inhibitors or serotonergic agonists/antagonists) during dynamic fPET imaging to directly test whether disrupting these gates alters a subject’s state-switching capacity or glucodynamic range. Additionally, investigating how these low-dimensional molecular gates break down in clinical cohorts will be important, such as in neurodegenerative disorders characterised by locus coeruleus degradation or neuropsychiatric illnesses marked by metabolic syndrome. Determining whether pathological "trapping" in sparsely connected metabolic states stems from a structural loss of these specific modulatory gates will validate this framework as a clinically viable biomarker for neuroenergetic health.

From a technical viewpoint, this work is the first to map ‘chronnectomic’ [17] or time-variant metabolic connectivity of the human brain. This represents an advancement for the metabolic connectivity field in the previous 5-years [105], moving from across-subject covariance approaches - which do not adequately reflect within-subject connectivity ([32, 106, 107]; see also [108]) - to the establishment of within-subject metabolic connectomes [32, 109, 110], to time-variant models (current results). Building upon foundational evidence about the known network state architecture of the brain, [29–31], we demonstrate that the metabolic organisation of the brain shows transitions between network states, and those dynamics are an important contributor to cognitive outcomes. It has been previously demonstrated that transient network state-switching is an important contributor to psychiatric [111] and neurodegenerative [112] illness, and that deterioration in the supply and use of cerebral glucose is a potential causative factor in many of these illnesses [113]. While speculative, this indicates that disruption of metabolic network state switching might ultimately prove to be an important biomarker for brain health, an application that is acknowledged to be difficult with fMRI given its known signal limitations [20, 114]. This work therefore represents an important next step in the understanding of the network dynamics of the fundamental metabolic properties of the brain, and future research should investigate contexts where deterioration of metabolic state switching may play a critical role in neuropathology, such as in metabolic syndrome, neurodegeneration and psychiatric conditions.

The results of this study should be considered in light of its strengths and limitations. A key strength is the use of fPET, allowing for the temporal characteristics of metabolic connectivity and signal dynamics to be studied. Another strength lies in examining the relationship between dynamic brain signals, metabolic network state switching and cognition across both younger and older adults. However, several limitations warrant consideration. First, the cross-sectional design precludes causal inference and limits our ability to assess trajectories of brain-cognition changes over time. Longitudinal studies are needed to establish whether the dynamic brain features identified here predict age-related cognitive *decline*, rather than age *differences*, as is the case here. Second, the 64 second duration used to compute the sliding window connectivity measures is relatively long and was necessitated by the 16 sec duration of the fPET frames. While sensitivity analysis at alternate window lengths supported the robustness and biological foundation of our results, future work will be strengthened by the use of shorter fPET frames of 3 secs (or less), which is just now becoming technically feasible [115]. Finally, this study focused on naturalistic resting-state dynamics, which capture intrinsic network organisation but do not index task-evoked brain activity - a valuable direction for future work. Our results should therefore be interpreted as reflecting the intrinsic metabolic architecture that supports endogenous and exogenous cognitive processing [116].

In conclusion, this study provides the first evidence that the brain’s metabolic network enters coherent states and undergoes dynamic state switching. Our results reframe the concept of macroscale metabolic brain dynamics from a purely abstract topological phenomenon to one explicitly bound by the real-time allocation of neuroenergetic substrates. The capacity to flexibly transition between network states is constrained by its underlying regional glucodynamic range and complexity, and maps to a specific neurochemical architecture. Our findings reveal that cognitive performance relies on a flexible metabolic landscape capable of network reconfigurations and sustained periods of associative system integration. In contrast, ageing perturbs this system, altering local glucodynamics and anchoring the brain within a sparsely connected baseline configuration at the expense of attaining states of associative and sensory system integration. Ultimately, these insights open up new avenues for research into the energetic costs of human cognition, providing a model for exploring how disrupted neuroenergetics and metabolic dysfunction compromise cognitive health in ageing and neurodegenerative disease.

## Materials and Methods

### Ethical Considerations

This study was approved by the Monash University Human Ethics Research Committee. The use of ionizing radiation was approved by the Monash Health Principal Medical Physicist Administration approved in accordance with the Australian Radiation Protection and Nuclear Safety Agency Code of Practice (2005). Participants provided informed consent to participate.

### Participants

The Monash MetConn simultaneous PET/MR dataset was used for the study. The dataset has been described in our previous work [11, 33]. Whole sample (N = 85) and younger (N = 40) and older (N = 45) age group differences in demographic characteristics and cognition are reported in detail in Supplement 2.1. Briefly, the mean age of the whole sample was 54.0 years (SD = 24.6), the younger group 27.8 years and the older group 75.5 years. The proportion of women in the younger group (60%) and older group (48%) was not significantly different. The older group had significantly greater mean systolic blood pressure (148mmHg vs 119mmHg), fasting blood glucose level (5.2 vs 4.7 mmol/L) and head motion (0.3mm vs 0.2mm).

### Data Acquisition

Full details of the cognitive tests are provided in Supplement 1.2. Briefly, participants completed the Hopkins Verbal Learning (HVLT) [117], task-switching [118], stop-signal [119] and digit symbol substitution [120] tests. Participants underwent a 90-minute simultaneous MR-PET scan in a Siemens (Erlangen, Germany) Biograph 3-Tesla molecular MR scanner. Half of the 260 MBq FDG tracer was administered as a bolus at commencement of the scan; and half was infused over 50 minutes at 36ml/hour. This protocol provides a balance between a rapid increase in signal-to-noise ratio and maintenance of signal-to-noise ratio over the scan period [44].

Participants were positioned supine in the scanner bore with their head in a 32-channel radiofrequency head coil. Non-functional T1 and T2 MRI scans were acquired in the first 12 minutes. The T1 3DMPRAGE scan parameters were: TA = 3.49 min, TR = 1640ms, TE = 2.34ms, flip angle = 8°, field of view = 256 × 256 mm^2^, voxel size = 1.0 × 1.0 × 1.0 mm^3^, 176 slices, sagittal acquisition. For the T2 FLAIR scan: TA = 5.52 min, TR = 5,000ms, TE = 396ms, field of view = 250 × 250 mm^2^, voxel size = .5 × .5 × 1 mm^3^, 160 slices). List-mode PET and T2* EPI BOLD-fMRI sequences began 13 minutes into the scan. The PET parameters were: voxel size = 1.39 x 1.39 x 5.0mm^3^; and the EPI BOLD-fMRI: TA = 40 minutes; TR = 1,000ms, TE = 39ms, FOV = 210 mm2, 2.4 × 2.4 × 2.4 mm^3^ voxels, 64 slices, ascending axial acquisition). The resting-state scan was undertaken while participants viewed a movie of a drone flying over the Hawaii Islands.

### MR & PET Data Preprocessing

Participants’ PET data were binned into 344 3D, 16s sinogram frames. Attenuation was corrected via the pseudo-CT method for hybrid PET-MR scanners[121] and 3D volumes constructed using the Ordinary Poisson-Ordered Subset Expectation Maximization algorithm with point spread function correction (3 iterations, 21 subsets). The reconstructed DICOM slices were converted to NIFTI format, 344 × 344 × 127 (size: 1.39 × 1.39 × 2.03 mm^3^). A single 4D NIFTI volume was constructed by concatenating the 3D volumes. The 4D PET volumes were motion corrected [122] and corrected for partial volume effects [123]. We used a 25% gray matter threshold [123] and surface-based spatial smoothing [124] with a Gaussian kernel with a full width at half maximum of 8 mm. To minimise the impact of the FDG baseline uptake on our dynamic connectivity measures, the fPET time-series were denoised using a standard anatomical component based noise correction method [125]. Previous work has shown this method to produce residual fPET timeseries reliably measuring the temporal dynamics of the glucose signal [11, 39, 47, 115].

### Time-Variant Network Connectivity

Time-variant metabolic connectivity was derived from connectivity matrices computed using sliding windows. Four consecutive frames (1:04 min window) with 50% overlap (32 seconds) were used, generating 150 windows per person. Following sliding-window extraction, all negative correlation coefficients were thresholded to a zero-bound floor. This approach permits the calculation of weighted graph metrics that assume non-negative edge weights and preserves the core topographic architecture of the time-varying networks [6, 126].

Time-resolved functional connectivity matrices were vectorized and clustered across the cohort using the k-means algorithm (k = 2-8, Cityblock/Manhattan distance, 5 random initialisations, 500 maximum iterations) to identify recurring dynamic connectivity states. Optimal model selection was evaluated using a combination of cluster validity criteria, specifically the Silhouette consensus profile and the within-cluster sum of squared errors (WCSS) elbow criterion. To assess whether the selected k = 4 cluster solution captured non-random dynamic structure, our empirically-derived clustering metrics were benchmarked against a null distribution generated via temporal block-bootstrap surrogates (n = 100). Timeseries were partitioned into non-overlapping blocks matched to the sliding-window length (w = 4 frames) and randomly shuffled prior to window extraction and clustering. This preserved local autocovariance structure while disrupting non-stationary dynamic transitions, allowing empirical WCSS and Silhouette scores to be directly evaluated against the surrogate baseline.

#### Sensitivity Analysis

The primary window length was selected to balance temporal resolution against the reliability of correlation estimation. Because the optimal temporal scale for dynamic metabolic connectivity remains unknown, additional sensitivity analyses were conducted using alternative window lengths (Supplement 2.7).

### Metabolic Network State Switching Measures

Temporal dynamics across these metabolic network states were quantified using three chronometric measures, as follows:

#### Total Transitions

The absolute frequency with which a participant shifted between any of the discrete network states over the course of the scan, serving as a primary index of global network flexibility.

#### Fractional Occupancy Rate

The percentage of windows spent in a given state by an individual participant.

#### Mean Dwell Time

The average number of consecutive windows a participant remained within a given state configuration during a single uninterrupted visit, capturing the continuous temporal stability and persistence of that specific network topology. Mean dwell time in windows (*W*) was converted to real-world duration in seconds using the formula: Duration = 64 + (*W* – 1) x 32, where 64 seconds represents the initial window length and 32 seconds the overlap-adjusted increment between successive windows.

### Graph Theoretical Metrics

To characterise the topological architecture of the metabolic network states, graph-theoretical metrics were calculated for each window and subsequently averaged across all windows assigned to the same network state. Local segregation was quantified using the weighted clustering coefficient, reflecting the tendency of neighbouring nodes to form densely interconnected communities [16]. Nodal strength centrality was calculated as the sum of all weighted connections attached to a given node [16]. Regions with high nodal strength were interpreted as highly connected hubs that contribute substantially to network integration.

### Glucose Cost of the Metabolic Network States

We computed a combined ‘Glucose Cost Index’ (GCI) score for each brain region by normalising clustering coefficients and centrality values within each state (min-max scaling to (0,1)) and taking their geometric mean. This approach identifies regions that have high local clustering (indicating specialised local processing) and high global centrality (indicating integration across the network). The geometric mean ensuring that regions scoring highly must possess at least moderate levels of both topological properties. For each subject and metabolic network state, we calculated the glucose GCI as the ratio of this combined clustering and centrality score to their regional CMR_GLC_. This metric captures metabolic cost - the amount of topological importance (both local and global) achieved per unit of glucose consumption. Higher GCI indicates a greater topological return (stronger clustering and/or centrality) for a given baseline glucose budget.

To assess the metabolic cost of the four network states, regions were ranked by their combined clustering and centrality score (geometric mean), and the top 25% were compared to the remaining 75% within each state using paired *t*-tests. To assess state-dependent differences in metabolic cost, we performed a repeated measures ANOVA on the GCI values of the top 25% regions across the four states, followed by Bonferroni-corrected post-hoc pairwise comparisons.

### Entropy and Standard Deviation of the Timeseries

We calculated regional sample entropy and the standard deviation of the FDG timeseries for each participant. We chose sample entropy as it has been well validated in neuroimaging studies, is sensitive to changes in the underlying time series, displays robustness to measurement noise and is adaptable for different time-series lengths and embedding dimensions [31]. Entropy was calculated per voxel using embedding dimension m=2 and tolerance r = 0.15 × SD of each individual time series. This relative tolerance ensures amplitude-invariance so that entropy reflects the temporal structure of the signal independent of its overall magnitude. The time series were not additionally standardized (i.e., z-scored) as the use of a relative tolerance renders such normalization unnecessary. The standard deviation of the FDG signal was also calculated voxel-wise to capture variability. We also chose not to standardise the timeseries values for SD calculation, as absolute fPET values carry physiologically relevant information (e.g., differences in metabolic rates) and standardisation can obscure these important effects. Moreover, subjects inherently have different variability in glucose use [39, 127], which standardisation would mask.

Regional entropy and standard deviation values were obtained by averaging voxel-wise values within each of the Schaefer 100 atlas regions [57]. A ‘glucodynamics index’ was also derived for analysis with network state switching measures, calculated as the average of z-scored entropy and SD measures across the 100 regions for each participant. While both measures are derived from the same time series, the use of a relative tolerance in entropy calculation (r = 0.15 × SD) ensures entropy reflects signal structure rather than amplitude, supporting the assertion that these measures capture different properties of the fPET time series.

### Head Motion

Relative head displacement was extracted from participants’ head motion correction reports as the mean head movement in each frame relative to the following frame. Framewise displacement values were calculated using translational and rotational head displacement between consecutive frames. Specifically, framewise displacement was calculated as the absolute sum of three translation and three rotation length derivates [128].

### Neurotransmitter profiles

To uncover the neurochemical profiles underlying the four metabolic states, we implemented an imaging receptor mapping framework leveraging the open-source *neuromaps* toolbox [68]. To bridge macroscale metabolic network topology and microscale chemoarchitecture, regional centrality of the 100 regions of the four states were spatially co-registered against an extensive in vivo repository of neurotransmitters receptors and transporters from high-resolution positron emission tomography (PET) and single-photon emission computed tomography (SPECT) maps acquired from healthy cohorts.

### Data Analyses

#### Metabolic Network State Switching and Cognition

Pearson correlation and multiple linear regression analyses were used to assess the associations between cognition and the metabolic network state switching measures (total state transitions, four state occupancy and mean dwell times, respectively). A single cognitive measure was derived from Principal Component Analysis of the four cognitive measures (HVLT, stop signal, category switch and digit symbol substitution).

#### Metabolic Network State Switching and Glucodynamic Signal Properties

Partial correlation analyses and multiple linear multiple regression analyses were used to assess the associations between the metabolic network state switching measures and the glucodynamics index, controlling head motion.

#### Metabolic Network States, Neurotransmitter Mapping and Pattern Decomposition

To characterise the neurochemical profile of the four metabolic network states, the *neuromaps* [68] repository of neurotransmitter receptors and transporters was used. Thirty-one independent neurotransmitter receptors and transporters were spatially correlated with centrality maps of the four network states (see Supplement Table S6 for list of transmitters). Statistical significance was assessed using permutation testing (n=1000 permutations), where the spatial arrangement of each state map was randomly shuffled to generate a null distribution of correlation coefficients. FDR correction was applied across all 124 metabolic network state-receptor pairs (p-FDR < 0.05).

Because individual neurotransmitter systems exhibit spatial overlap and collinearity due to shared cellular distribution profiles and cortical hierarchies [68], we also performed an unsupervised pattern decomposition via Principal Component Analysis (PCA) to identify latent neurochemical axes that co-vary with the metabolic state topographies. To construct the input matrix, we first computed the spatial correlation between each of the 31 receptor maps and each of the four metabolic state maps (i.e., a 31 × 4 matrix of correlation coefficients; receptors as rows, states as columns). Each column was z-score normalised to achieve zero mean and unit variance prior to PCA. This ensures that variance in the PCA is driven by differential receptor profiles across states rather than by differences in overall correlation strength. We note, however, that with only four state columns, the PCA decomposes a limited amount of variance; and focus interpretation mostly on consistent patterns. The per-receptor projections onto each PC are referred to as scores (or receptor weights), and we report these for the top-contributing receptors. The loadings, i.e., the correlation between each PC and the original state variables - were also computed to characterise how each PC relates to the four metabolic state topographies.

#### Age Group Differences in Metabolic Network State Switching and Glucodynamic Signal Properties

A series of general linear models (GLMs) was run to test for age group differences in fPET standard deviation and entropy at the region level, with head motion as a covariate. Partial eta squared (η^2^_p_) was used to quantify the age effect sizes. Age group comparisons of the metabolic network state switching measures were undertaken with t-tests.

#### False Discovery Rate Correction and Outlier Detection

Across all correlational and inferential testing, individual data points meeting the a priori exclusion criteria of exceeding three standard deviations from the cohort mean on any given variable were omitted from that specific analysis via pairwise deletion. This approach maximised statistical power by retaining subjects with valid data points across alternative metrics, with n-sizes reported for each distinct statistical calculation. Finally, to control for the inflation of Type I errors inherent to multiple concurrent testing spaces, the Benjamini-Hochberg False Discovery Rate (FDR) correction procedure was applied for each independent block of contrasts (e.g., cognition PC × network state metrics; region-level age GLMs of fPET SD) with statistical significance established at a corrected threshold of p-FDR < 0.05

## Data and Code Availability Statement

All statistics supporting the results of this study are given in the Supplementary Information. The datasets and code used for the study are available https://osf.io/nprbs

## Author Contributions

SDJ and GFE conceived the project. HD and SDJ, CM and GFE designed and developed the manuscript. HD and EL analysed the data. HD wrote the manuscript, SDJ, GFE and CM reviewed/edited the manuscript. All authors read and approved the final manuscript.

## Acknowledgements

This work was supported by Australian Research Council (ARC) Discovery Project DP25010302 and ARC Fellowship FT250100206.

We thank Robert Di Paolo, Gerard Murray, M. Navyaan Siddiqui, Katharina Voigt, Richard McIntyre, Lauren Hudswell and the staff at Monash Biomedical Imaging for their contributions to data acquisition and image reconstruction.

