## Supplement for "Metabolic brain networks switch between a sparsely connected baseline and highly integrated states to support cognition"

~ Supplementary Methods and Results ~

**1. Supplementary Methods** **2**
1.1 Participants 2
1.2 Cognitive Tests 2

**2. Supplementary Results** **3**
2.1 Sample Characteristics 3
2.2 Cluster Number Validation 4
2.3 Regional profiles of four dynamic metabolic states 5
2.4 Timeseries Standard Deviation and Sample Entropy 9
2.5 Neurochemical system alignment with metabolic network states 10
2.6 Region Glucodynamics 11
2.7 Sensitivity Analysis 14

### 1. Supplementary Methods

##### *1.1 Participants*

Local advertising was used to recruit 90 participants from the general community. A screening interview was conducted to ensure that participants had the capacity to provide informed consent. Participants were also screened to ensure that they did not have a diagnosis of diabetes, neurological or psychiatric illness, claustrophobia or non-MR compatible implants. They were also excluded if they had received a clinical or research PET scan in the past 12 months. Women were screened for current or suspected pregnancy. Participants received a $100 voucher for participating in the study. Five participants were excluded from further analyses due to excessive head motion (N=2), incomplete PET scan or image reconstruction (N=2) and out of range data (N = 1).

##### *1.2 Cognitive Tests*

**Hopkins Verbal Learning Test (HVLT).** The HVLT is a three-trial list learning and free recall task. The learning trials comprised 12 words, four words from each of three semantic categories [1]. Approximately 20–25 minutes after the learning trials, participants completed delayed recall and recognition trials. The delayed recall required free recall of any of the 12 words. The recognition trial comprised 24 words, including the 12 target words and 12 false-positives, six semantically related, and six semantically unrelated words. Delayed recall was calculated as the total words recalled.

**Task Switching.** For task switching, a computerised test was used in which participants were presented with a word and had to perform a categorisation task. The categorisation task was dependant on two cues that appeared across the trials. One cue was a heart symbol, for which participants were asked to categorise the word presented via a key press as either a LIVING or a NON-LIVING object. If the cue was an arrow-cross, participants were asked to categorise the word as either BIGGER or SMALLER than a basketball. The cue was randomised across trials. Half the trials were switch trials and half were non-switch trials. Half the switch and non-switch trials was congruent in the key presses for either task, half was incongruent. The task switching measure was latency of correctly responding to a switch trial [2].

**Stop Signal.** The stop signal trial was a computer-based test [3]. Participants were required to press the left response key if an arrow on screen pointed left and the right response key if the arrow pointed right. If a signal beep sounded, participants were instructed stop their response. The delay between presentation of an arrow and signal beep started at 250ms and was altered up or down by 50ms based on performance. The delay increased up to 1150ms if the previous stop signal trial was successful and decreased down to 50ms if the previous stop signal trial was unsuccessful. The stimulus onset asynchrony between the onset of a fixation circles at the start of each trial was 2000ms. Reaction time in the stop signal trials was recorded.

**Digit Symbol Substitution.** A computer-based task presenting participants with a matric of 18 column and 16 rows [4]. Participants were required to translate symbols shown above the matrix in a key into digits in the matrix. The trial lasted two minutes. Performance was measured as total count of correct responses.

#### 2. Supplementary Results

##### *2.1 Sample Characteristics*

The characteristics of the whole sample (N =85), as well as the younger (N = 40) and older (N = 45) participants, are shown in Supplementary Table S1. The mean age of the whole sample was 53.3 years (SD = 24.8). The proportion of women was 53%. Average BMI was 25.0 kg/m$^2$, resting heart rate was 79 BPM and systolic and diastolic blood pressure were 135 and 82 mmHg, respectively. The mean fasting blood glucose was 5.0 mmol/L.

The mean age of the younger group was 27.9 years and the older group 75.8 years. The proportion of women in the younger group (55%) and older group (51%) was not significantly different. The average years of education was higher in the younger (18.0) than the older group (16.2). The older group had significantly higher mean systolic blood pressure than the younger group (150mmHg vs 119mmHg). The older group also had a higher fasting blood glucose level (5.2 vs 4.8 mmol/L). A total of 27 people in the older group and eight in the younger group met diagnostic guidelines for hypertension. These rates are on par with those in the Australian adult population [5]. Although older people had a slightly higher mean BMI than younger people (25.7 vs 24.1 kg/m$^2$), the difference was not statistically significant. Nine participants (12%) would meet the definition for obesity (BMI above 30), including three younger and six older participants. This is below the 32% of the Australian adult population that is obese [5]. Older adults performed worse on all cognitive tests than younger adults.

**Table S1. Demographics for the whole sample and comparison of older and younger groups.** Continuous variables are mean (standard deviation); categorical variables are number and percentage.

| | Whole sample (N = 85) | | Younger (N = 40) | | Older (N = 45) | | Younger vs Older p-value[1] |
|---|---|---|---|---|---|---|---|
| | Mean | SD | Mean | SD | Mean | SD | |
| Age | 53.3 | 24.8 | 27.9 | 6.2 | 75.8 | 6.1 | NA |
| Sex (number (%) females) | 45 (53%) | | 22 (55%) | | 23 (51%) | | 0.820 |
| Education (years) | 17.0 | 3.6 | 18.0 | 2.7 | 16.2 | 4.0 | 0.026 |
| Glucose (mmol/L) | 5.0 | 0.5 | 4.8 | 0.4 | 5.2 | 0.6 | 0.000 |
| Systolic BP (mmHG) | 135.4 | 26.4 | 119.0 | 17.2 | 149.9 | 24.5 | 0.000 |
| Diastolic BP (mmHG) | 81.6 | 12.9 | 78.5 | 13.1 | 84.4 | 12.1 | 0.032 |
| Resting HR (BPM) | 78.5 | 15.4 | 82.5 | 16.9 | 75.0 | 13.1 | 0.025 |
| BMI (kg/m²) | 25.0 | 4.1 | 24.1 | 4.6 | 25.7 | 3.6 | 0.080 |
| HVLT delayed recall | 8.2 | 2.7 | 9.3 | 2.5 | 7.2 | 2.6 | 0.000 |
| Category switch RT | 1.80 | 0.65 | 1.42 | 0.40 | 2.13 | 0.65 | 0.000 |
| Digit symbol sub (correct) | 44.3 | 23.5 | 63.6 | 16.5 | 27.6 | 13.9 | 0.000 |
| Stop signal RT | 0.57 | 0.13 | 0.53 | 0.13 | 0.60 | 0.12 | 0.011 |

[1]P-values are based on T-test for continuous (2-sided) and Ch-square for categorical variables. Note: Education was not available for two younger and one older adult. One older participant's digit symbol substitution score was more than 3 standard deviations below the mean, and their data was excluded from analyses of cognition.

##### *2.2 Cluster Number Validation*

Clustering was evaluated across a range of solutions (k = 2 - 8) using the elbow criterion alongside the mean silhouette consensus profile (Fig S1). The WCSS evaluation demonstrated subtle changes across cluster number, while silhouette coefficients remained positive at k = 4 prior to degrading toward zero at higher cluster numbers (k > 6). Combined with prior empirical literature (see main manuscript), we selected k = 4 as the optimal model solution. Block-bootstrap surrogate testing (n = 20) confirmed that the k = 4 model captures non-random dynamic structure, yielding a significantly lower WCSS in real data compared to phase-shuffled surrogates (p = 0.009; Cohen's d = -71.7). Thus, k = 4 strikes a balance between data-driven cluster separation, parsimony, and neurophysiological interpretability, while reflecting that connectivity states likely occupy a continuous topological space rather than purely isolated clusters

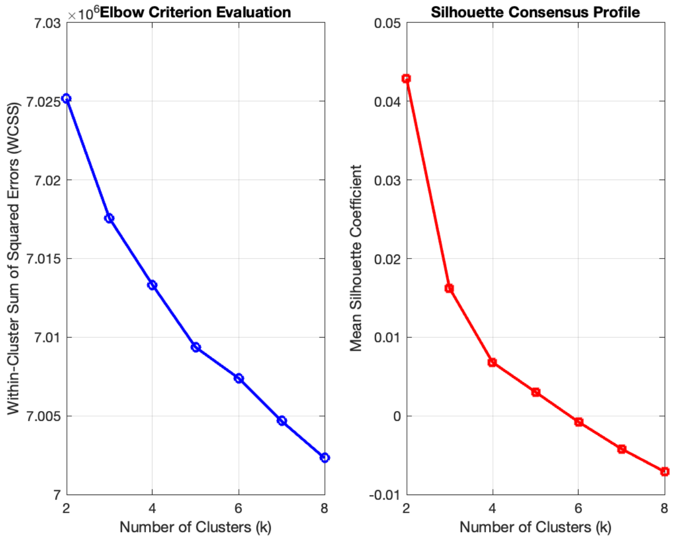


**Fig S1. Cluster validation profiles for the selection of the optimal dynamic metabolic network state model.** Evaluation of number of clusters across a range of cluster solutions (k = 2 to 8) derived from sliding-window functional PET (fPET) connectivity matrices (see Methods, main manuscript).

##### *2.3 Regional profiles of four dynamic metabolic states*

**Table S2. Dynamic metabolic State 1.** Rank order of regional clustering and centrality for State 1. Blue shading is top 25% of regions.

| State 1: Clustering | | State1: Centrality | |
|---|---|---|---|
| SVA B: Medial Posterior Prefrontal 1 R | 0.199 | CON B: Lateral Prefrontal Cortexv 1 R | 12.8 |
| DEF C: Parahippocampal Cortex 1 R | 0.135 | DEF B: Ventral Prefrontal Cortex 2 L | 11.1 |
| DEF A: Dorsal Prefrontal Cortex 1 L | 0.130 | CON A: Lateral Prefrontal Cortex 1 L | 11.1 |
| SVA B: Medial Posterior Prefrontal 1 L | 0.125 | CON B: Temp 1 R | 10.2 |
| TP 1 R | 0.125 | VIS Cent: Extra Striate Cortex 2 L | 10.2 |
| DEF B: Ventral Prefrontal Cortex 1 R | 0.125 | VIS Cent: Extra Striate Cortex 2 R | 10.0 |
| DA B: Frontal Eye Fields 1 R | 0.123 | CON B: Lateral Prefrontal Cortexv 1 L | 9.8 |
| DEF B: Ventral Prefrontal Cortex 1 L | 0.123 | VIS Cent: Extra Striate Cortex 3 L | 9.6 |
| SOM A: 3 R | 0.122 | SOM B: Auditory 1 L | 9.1 |
| VIS Perhiph: Extra Striate Superior 1 R | 0.121 | DA A: TEMP Occipital 1 R | 9.1 |
| VIS Perhiph: Extra Striate Inferior 1 R | 0.119 | LIM B: Orbital Frontal Cortex 0 R | 8.9 |
| DA B: Frontal Eye Fields 1 L | 0.117 | SOM B: Auditory 1 R | 8.9 |
| DEF B: Temp 2 L | 0.116 | LIM A Temp Pole 1 L | 8.7 |
| CON A: Lateral Prefrontal Cortex 2 L | 0.116 | DEF B: Inferior Parietal Lobule 1 L | 8.6 |
| DEF A: Medial Prefrontal Cortex 1 L | 0.116 | DEF B: Dorsal Prefrontal Cortex 1 L | 8.4 |
| SVA A: Insula: 1 R | 0.115 | CON A: Lateral Prefrontal Cortex 1 R | 8.2 |
| TP 3 R | 0.113 | DA A: Parietal Occipital 1 R | 8.1 |
| VIS Cent: Extra Striate Cortex 1 L | 0.112 | DA A: TEMP Occipital 1 L | 8.1 |
| LIM B: Orbital Frontal Cortex 1 L | 0.112 | DEF B: Ventral Prefrontal Cortex 2 R | 7.5 |
| TP 1 L | 0.112 | LIM B: Orbital Frontal Cortex 1 L | 7.3 |
| DEF C: RetroSuperior Parietal Lobuleenial 1 R | 0.111 | DEF A: Medial Prefrontal Cortex 1 R | 7.1 |
| VIS Cent: Extra Striate Cortex 1 R | 0.111 | VIS Cent: Extra Striate Cortex 3 R | 7.0 |
| SVA A: Insula: 2 L | 0.111 | SVA B: Lateral Prefrontal Cortex 1 R | 7.0 |
| VIS Perhiph: Extra Striate CortexSup 1 L | 0.110 | DEF B: Dorsal Prefrontal Cortex 1 R | 6.8 |
| LIM A: Temp Pole 1 R | 0.110 | TP 2 R | 6.7 |
| DEF B: Ventral Prefrontal Cortex 2 R | 0.109 | VIS Cent: Extra Striate Cortex 1 R | 6.7 |
| CON A: Lateral Prefrontal Cortex 2 R | 0.109 | VIS Perhiph: Striate Cortex Calcarine 1 L | 6.6 |
| DEF B: Temp 1 L | 0.109 | SOM B: S2 2 L | 6.5 |
| DA A: Parietal Occipital 1 L | 0.109 | DEF A: Inferior Parietal Lobule 1 R | 6.2 |
| DEF B: Lateral Prefrontal Cortex 1 L | 0.109 | CON B: inferior parietal lobule 1 R | 6.2 |
| DA A: Superior Parietal Lobule 1 L | 0.108 | DEF B: Temp 1 L | 6.1 |
| SVA B: Lateral Prefrontal Cortex 1 L | 0.108 | DEF A: Precuneus Posterior Cingulate Cortex1 L | 5.9 |
| CON A: Intraparietal Sulcus 1 L | 0.107 | VIS Cent: Striate Cortex 1 L | 5.6 |
| VIS Perhiph: Extra Striate Inferior 1 L | 0.107 | VIS Perhiph: Extra Striate CortexSup 1 L | 5.6 |
| DEF A: Medial Prefrontal Cortex 1 R | 0.107 | CON B: Lateral Prefrontal Cortexd 1 R | 5.5 |
| VIS Perhiph: Striate Cortex Calcarine 1 R | 0.104 | VIS Perhiph: Striate Cortex Calcarine 1 R | 5.4 |
| CON C: Precuneus 1 L | 0.104 | VIS Cent: Extra Striate Cortex 1 L | 5.3 |
| LIM A: Temp Pole 2 L | 0.104 | DEF B: Temp 2 L | 5.2 |
| SVA B: Lateral Prefrontal Cortex 1 R | 0.104 | DA A: Parietal Occipital 1 L | 5.1 |
| DA A: Superior Parietal Lobule 1 R | 0.103 | DEF B: Ventral Prefrontal Cortex 1 R | 5.1 |
| SVA A: Parietal Operculum 1 L | 0.103 | SVA B: Lateral Prefrontal Cortex 1 L | 5.1 |
| DEF A: Precuneus Posterior Cingulate Cortex1 L | 0.103 | DEF A: Medial Prefrontal Cortex 1 L | 5.0 |
| SOM B: S2 1 R | 0.103 | TP 1 L | 4.9 |
| VIS Cent: Extra Striate Cortex 2 R | 0.102 | CON A: Lateral Prefrontal Cortex 2 R | 4.9 |
| DEF A: Precuneus Posterior Cingulate Cortex 1 R | 0.102 | VIS Perhiph: Extra Striate Inferior 1 R | 4.9 |
| VIS Perhiph: Striate Cortex Calcarine 1 L | 0.102 | SVA A: Insula: 1 R | 4.8 |
| LIM A Temp Pole 1 L | 0.102 | LIM A: Temp Pole 1 R | 4.6 |
| DEF C: RetroSuperior Parietal Lobuleenial 1 L | 0.102 | CON C: Precuneus 1 R | 4.4 |
| SOM B: S2 1 L | 0.102 | DEF C: RetroSuperior Parietal Lobuleenial 1 L | 4.3 |
| SVA B: Inferior Parietal Lobule 1 R | 0.100 | TP 3 R | 4.0 |
| CON C: Cingulate Posterior 1 L | 0.100 | CON C: Precuneus 1 L | 3.9 |
| DEF B: Inferior Parietal Lobule 1 L | 0.100 | SVA A: Insula: 2 L | 3.6 |
| CON B: Temp 1 R | 0.100 | VIS Perhiph: Extra Striate Inferior 1 L | 3.6 |
| VIS Cent: Striate Cortex 1 L | 0.099 | CON A: Intraparietal Sulcus 1 L | 3.3 |
| VIS Cent: Extra Striate Cortex 3 R | 0.098 | VIS Perhiph: Extra Striate Superior 1 R | 3.2 |
| CON B: Lateral Prefrontal Cortexd 1 R | 0.097 | SVA A: Parietal Operculum 1 L | 3.1 |
| DEF A: Inferior Parietal Lobule 1 R | 0.097 | DEF B: Ventral Prefrontal Cortex 1 L | 3.1 |
| CON C: Precuneus 1 R | 0.096 | SOM B: S2 1 L | 2.8 |
| LIM B: Orbital Frontal Cortex 0 R | 0.096 | DEF A: Dorsal Prefrontal Cortex 1 R | 2.7 |
| DEF A: Dorsal Prefrontal Cortex 1 R | 0.096 | SVA A: Parietal Operculum 1 R | 2.6 |
| CON B: inferior parietal lobule 1 R | 0.096 | DA A: Superior Parietal Lobule 1 L | 2.5 |
| DA A: Parietal Occipital 1 R | 0.095 | CON A: Lateral Prefrontal Cortex 2 L | 2.4 |
| CON C: Precuneus 2 L | 0.093 | SOM B: S2 2 R | 2.4 |
| DA A: TEMP Occipital 1 L | 0.093 | LIM A: Temp Pole 2 L | 2.3 |
| DA A: TEMP Occipital 1 R | 0.093 | SOM B: Cent 1 L | 2.3 |
| TP 2 R | 0.091 | DEF B: Lateral Prefrontal Cortex 1 L | 2.2 |
| VIS Cent: Extra Striate Cortex 3 L | 0.091 | TP 1 R | 2.2 |
| SVA A: Frontal Medial 1 L | 0.090 | DA A: Superior Parietal Lobule 1 R | 2.2 |
| VIS Cent: Extra Striate Cortex 2 L | 0.090 | DEF C: RetroSuperior Parietal Lobuleenial 1 R | 2.1 |
| CON A: Lateral Prefrontal Cortex 1 R | 0.090 | DEF A: Precuneus Posterior Cingulate Cortex 1 R | 2.0 |
| SOM B: S2 2 L | 0.089 | SOM B: S2 1 R | 1.8 |
| SOM B: Auditory 1 L | 0.088 | CON A: Intraparietal Sulcus 1 R | 1.6 |
| SOM B: Auditory 1 R | 0.088 | CON C: Precuneus 2 L | 1.5 |
| DEF B: Dorsal Prefrontal Cortex 1 R | 0.087 | DEF C: Parahippocampal Cortex 1 R | 1.5 |
| SOM B: Cent 1 L | 0.085 | SVA B: Medial Posterior Prefrontal 1 L | 1.4 |
| DEF C: Parahippocampal Cortex 1 L | 0.085 | SVA B: Inferior Parietal Lobule 1 R | 1.2 |
| SVA A: Parietal Operculum 1 R | 0.085 | CON C: Cingulate Posterior 1 L | 1.1 |
| CON B: Lateral Prefrontal Cortexv 1 R | 0.084 | SOM B: Cent 1 R | 0.9 |
| DEF B: Dorsal Prefrontal Cortex 1 L | 0.084 | DA B: Post Cent 1 L | 0.9 |
| CON B: Lateral Prefrontal Cortexv 1 L | 0.083 | SVA A: Parietal Medial 1 L | 0.9 |
| DEF B: Ventral Prefrontal Cortex 2 L | 0.082 | SOM A: 2 L | 0.8 |
| SOM B: S2 2 R | 0.081 | SOM A: 1 L | 0.7 |
| DA B: Post Cent 1 L | 0.077 | CON C: Cingulate Posterior 1 R | 0.7 |
| CON A: Lateral Prefrontal Cortex 1 L | 0.074 | DEF A: Dorsal Prefrontal Cortex 1 L | 0.7 |
| SVA A: Parietal Medial 1 L | 0.068 | DA B: Frontal Eye Fields 1 R | 0.6 |
| SVA A: Frontal Medial 1 R | 0.066 | SVA A: Insula: 1 L | 0.6 |
| CON A: Intraparietal Sulcus 1 R | 0.057 | SVA A: Frontal Medial 1 R | 0.6 |
| SVA A: Insula: 1 L | 0.054 | DA B: Post Cent 2 R | 0.6 |
| SOM B: Cent 1 R | 0.053 | DA B: Frontal Eye Fields 1 L | 0.5 |
| DA B: Post Cent 2 R | 0.043 | SVA B: Medial Posterior Prefrontal 1 R | 0.5 |
| DA B: Post Cent 1 R | 0.041 | SOM A: 4 R | 0.5 |
| SOM A: 2 L | 0.033 | SVA A: Frontal Medial 1 L | 0.5 |
| CON C: Cingulate Posterior 1 R | 0.023 | DA B: Post Cent 2 L | 0.5 |
| SOM A: 1 L | 0.013 | SVA A: Parietal Medial 1 R | 0.5 |
| DA B: Post Cent 2 L | 0.000 | SOM A: 1 R | 0.4 |
| DA B: Post Cent 3 L | 0.000 | DEF C: Parahippocampal Cortex 1 L | 0.4 |
| SOM A: 1 R | 0.000 | DA B: Post Cent 1 R | 0.4 |
| SOM A: 2 R | 0.000 | DA B: Post Cent 3 L | 0.3 |
| SOM A: 4 R | 0.000 | SOM A: 3 R | 0.3 |
| SVA A: Parietal Medial 1 R | 0.000 | SOM A: 2 R | 0.2 |

**Table S3. Dynamic metabolic State 2.** Rank order of regional clustering and centrality for State 2. Blue shading is top 25% of regions.

| State 2: Clustering | | State 2: Centrality | |
|---|---|---|---|
| CON A: Intraparietal Sulcus 1 R | 0.129 | DEF B: Dorsal Prefrontal Cortex 1 R | 12.4 |
| SVA A: Parietal Medial 1 L | 0.129 | CON B: Lateral Prefrontal Cortexv 1 R | 12.0 |
| SVA A: Parietal Medial 1 R | 0.127 | SVA B: Lateral Prefrontal Cortex 1 R | 11.1 |
| CON C: Cingulate Posterior 1 L | 0.126 | CON B: Lateral Prefrontal Cortexd 1 R | 10.9 |
| SOM A: 1 R | 0.125 | CON A: Lateral Prefrontal Cortex 1 R | 10.6 |
| DEF B: Dorsal Prefrontal Cortex 1 L | 0.124 | DA B: Frontal Eye Fields 1 L | 10.3 |
| DA B: Post Cent 2 R | 0.123 | DEF B: Dorsal Prefrontal Cortex 1 L | 10.2 |
| DEF B: Lateral Prefrontal Cortex 1 L | 0.122 | DEF A: Dorsal Prefrontal Cortex 1 R | 10.0 |
| CON A: Lateral Prefrontal Cortex 2 L | 0.122 | DA B: Frontal Eye Fields 1 R | 9.9 |
| CON B: Lateral Prefrontal Cortexd 1 R | 0.121 | SOM A: 1 L | 9.5 |
| SVA B: Lateral Prefrontal Cortex 1 L | 0.120 | SOM A: 2 L | 9.4 |
| SOM A: 2 R | 0.120 | CON A: Lateral Prefrontal Cortex 2 R | 9.2 |
| DA B: Post Cent 3 L | 0.120 | CON A: Lateral Prefrontal Cortex 1 L | 8.9 |
| SOM A: 4 R | 0.120 | SVA B: Lateral Prefrontal Cortex 1 L | 8.5 |
| DEF B: Dorsal Prefrontal Cortex 1 R | 0.119 | SOM A: 4 R | 8.2 |
| SVA A: Frontal Medial 1 R | 0.119 | DA A: Superior Parietal Lobule 1 L | 8.2 |
| SOM A: 3 R | 0.118 | SVA A: Frontal Medial 1 R | 8.1 |
| LIM A Temp Pole 1 L | 0.118 | SVA B: Medial Posterior Prefrontal 1 R | 8.1 |
| CON C: Cingulate Posterior 1 R | 0.117 | CON A: Intraparietal Sulcus 1 L | 8.0 |
| SOM B: S2 2 R | 0.117 | DEF A: Dorsal Prefrontal Cortex 1 L | 7.9 |
| SOM B: S2 1 R | 0.117 | SVA A: Frontal Medial 1 L | 7.8 |
| DA B: Post Cent 1 R | 0.117 | DEF B: Ventral Prefrontal Cortex 2 R | 7.7 |
| DEF A: Medial Prefrontal Cortex 1 L | 0.117 | DEF B: Lateral Prefrontal Cortex 1 L | 7.4 |
| CON A: Lateral Prefrontal Cortex 2 R | 0.117 | CON B: inferior parietal lobule 1 R | 7.3 |
| CON A: Lateral Prefrontal Cortex 1 L | 0.117 | CON B: Lateral Prefrontal Cortexv 1 L | 7.3 |
| DA B: Frontal Eye Fields 1 R | 0.117 | DA A: Superior Parietal Lobule 1 R | 7.2 |
| DEF B: Ventral Prefrontal Cortex 1 R | 0.116 | SOM A: 2 R | 7.0 |
| SVA A: Frontal Medial 1 L | 0.116 | DA B: Post Cent 2 R | 6.7 |
| CON C: Precuneus 1 R | 0.116 | CON C: Precuneus 1 R | 6.6 |
| SVA B: Medial Posterior Prefrontal 1 R | 0.116 | DA B: Post Cent 3 L | 6.4 |
| SOM A: 2 L | 0.115 | DEF B: Ventral Prefrontal Cortex 2 L | 6.4 |
| DA B: Post Cent 2 L | 0.114 | SVA B: Medial Posterior Prefrontal 1 L | 6.1 |
| SVA B: Inferior Parietal Lobule 1 R | 0.114 | DA B: Post Cent 1 R | 6.1 |
| CON A: Intraparietal Sulcus 1 L | 0.113 | CON C: Cingulate Posterior 1 R | 5.8 |
| DEF A: Precuneus Posterior Cingulate Cortex 1 R | 0.113 | SVA A: Parietal Medial 1 R | 5.8 |
| CON C: Precuneus 2 L | 0.113 | DEF A: Medial Prefrontal Cortex 1 R | 5.7 |
| DEF A: Dorsal Prefrontal Cortex 1 L | 0.112 | DEF A: Precuneus Posterior Cingulate Cortex1 L | 5.7 |
| SVA B: Lateral Prefrontal Cortex 1 R | 0.112 | CON A: Intraparietal Sulcus 1 R | 5.6 |
| DEF A: Medial Prefrontal Cortex 1 R | 0.111 | SOM B: Cent 1 R | 5.5 |
| DEF A: Dorsal Prefrontal Cortex 1 R | 0.111 | DEF B: Inferior Parietal Lobule 1 L | 5.3 |
| CON A: Lateral Prefrontal Cortex 1 R | 0.111 | SOM A: 1 R | 5.3 |
| SVA A: Parietal Operculum 1 R | 0.110 | SOM B: Cent 1 L | 5.2 |
| SVA B: Medial Posterior Prefrontal 1 L | 0.110 | CON C: Precuneus 2 L | 5.1 |
| SOM B: Cent 1 R | 0.108 | CON A: Lateral Prefrontal Cortex 2 L | 5.0 |
| DA B: Post Cent 1 L | 0.108 | DEF A: Precuneus Posterior Cingulate Cortex 1 R | 4.8 |
| CON B: Lateral Prefrontal Cortexv 1 R | 0.108 | SVA A: Parietal Operculum 1 R | 4.8 |
| DEF A: Precuneus Posterior Cingulate Cortex1 L | 0.107 | SVA B: Inferior Parietal Lobule 1 R | 4.6 |
| SOM B: Auditory 1 R | 0.107 | DA B: Post Cent 2 L | 4.2 |
| LIM B: Orbital Frontal Cortex 0 R | 0.107 | SOM B: Auditory 1 R | 4.1 |
| SOM B: Cent 1 L | 0.106 | SVA A: Parietal Medial 1 L | 4.1 |
| CON B: Lateral Prefrontal Cortexv 1 L | 0.106 | SOM B: S2 2 R | 4.0 |
| DEF B: Ventral Prefrontal Cortex 2 R | 0.105 | SOM A: 3 R | 3.7 |
| SOM A: 1 L | 0.105 | DEF B: Ventral Prefrontal Cortex 1 R | 3.6 |
| CON B: inferior parietal lobule 1 R | 0.105 | DEF A: Medial Prefrontal Cortex 1 L | 3.5 |
| DA B: Frontal Eye Fields 1 L | 0.104 | CON B: Temp 1 R | 3.4 |
| DEF B: Temp 1 L | 0.104 | SVA A: Insula: 1 R | 3.1 |
| DA A: Superior Parietal Lobule 1 R | 0.103 | LIM B: Orbital Frontal Cortex 0 R | 3.1 |
| DA A: Superior Parietal Lobule 1 L | 0.103 | CON C: Cingulate Posterior 1 L | 3.0 |
| VIS Cent: Extra Striate Cortex 3 R | 0.103 | DEF A: Inferior Parietal Lobule 1 R | 2.7 |
| DEF B: Inferior Parietal Lobule 1 L | 0.102 | CON C: Precuneus 1 L | 2.1 |
| SVA A: Insula: 1 R | 0.101 | SVA A: Parietal Operculum 1 L | 2.0 |
| DEF B: Ventral Prefrontal Cortex 2 L | 0.099 | DA B: Post Cent 1 L | 1.9 |
| CON B: Temp 1 R | 0.098 | VIS Cent: Extra Striate Cortex 3 R | 1.5 |
| CON C: Precuneus 1 L | 0.097 | VIS Perhiph: Extra Striate Superior 1 R | 1.5 |
| DEF A: Inferior Parietal Lobule 1 R | 0.096 | DA A: Parietal Occipital 1 R | 1.4 |
| DA A: Parietal Occipital 1 R | 0.094 | SOM B: Auditory 1 L | 1.3 |
| SVA A: Parietal Operculum 1 L | 0.090 | TP 2 R | 1.1 |
| TP 2 R | 0.084 | LIM A: Temp Pole 1 R | 0.9 |
| DEF C: RetroSuperior Parietal Lobuleenial 1 R | 0.078 | LIM A Temp Pole 1 L | 0.9 |
| DEF B: Ventral Prefrontal Cortex 1 L | 0.077 | SOM B: S2 1 R | 0.8 |
| VIS Perhiph: Striate Cortex Calcarine 1 R | 0.076 | VIS Perhiph: Striate Cortex Calcarine 1 L | 0.8 |
| VIS Perhiph: Extra Striate Superior 1 R | 0.069 | DEF C: RetroSuperior Parietal Lobuleenial 1 L | 0.8 |
| DEF C: RetroSuperior Parietal Lobuleenial 1 L | 0.060 | LIM B: Orbital Frontal Cortex 1 L | 0.7 |
| LIM A: Temp Pole 1 R | 0.058 | DEF C: RetroSuperior Parietal Lobuleenial 1 R | 0.7 |
| VIS Perhiph: Striate Cortex Calcarine 1 L | 0.054 | DEF B: Ventral Prefrontal Cortex 1 L | 0.7 |
| VIS Perhiph: Extra Striate CortexSup 1 L | 0.052 | VIS Perhiph: Extra Striate CortexSup 1 L | 0.7 |
| VIS Cent: Striate Cortex 1 L | 0.051 | VIS Perhiph: Extra Striate Inferior 1 L | 0.6 |
| SOM B: Auditory 1 L | 0.050 | SVA A: Insula: 2 L | 0.5 |
| TP 1 R | 0.047 | SOM B: S2 2 L | 0.5 |
| SOM B: S2 1 L | 0.040 | VIS Perhiph: Extra Striate Inferior 1 R | 0.5 |
| VIS Perhiph: Extra Striate Inferior 1 L | 0.025 | VIS Cent: Extra Striate Cortex 2 R | 0.5 |
| SVA A: Insula: 2 L | 0.024 | VIS Cent: Extra Striate Cortex 3 L | 0.4 |
| LIM B: Orbital Frontal Cortex 1 L | 0.021 | TP 1 R | 0.4 |
| SOM B: S2 2 L | 0.020 | VIS Cent: Extra Striate Cortex 2 L | 0.4 |
| VIS Perhiph: Extra Striate Inferior 1 R | 0.020 | VIS Cent: Striate Cortex 1 L | 0.4 |
| VIS Cent: Extra Striate Cortex 2 R | 0.018 | VIS Perhiph: Striate Cortex Calcarine 1 R | 0.4 |
| VIS Cent: Extra Striate Cortex 1 L | 0.000 | SOM B: S2 1 L | 0.4 |
| VIS Cent: Extra Striate Cortex 2 L | 0.000 | VIS Cent: Extra Striate Cortex 1 R | 0.4 |
| VIS Cent: Extra Striate Cortex 3 L | 0.000 | TP 3 R | 0.3 |
| DA A: TEMP Occipital 1 L | 0.000 | DA A: TEMP Occipital 1 L | 0.3 |
| DA A: Parietal Occipital 1 L | 0.000 | VIS Cent: Extra Striate Cortex 1 L | 0.3 |
| SVA A: Insula: 1 L | 0.000 | DA A: Parietal Occipital 1 L | 0.3 |
| LIM A: Temp Pole 2 L | 0.000 | DA A: TEMP Occipital 1 R | 0.3 |
| DEF B: Temp 2 L | 0.000 | SVA A: Insula: 1 L | 0.3 |
| DEF C: Parahippocampal Cortex 1 L | 0.000 | DEF B: Temp 2 L | 0.3 |
| TP 1 L | 0.000 | DEF B: Temp 1 L | 0.2 |
| VIS Cent: Extra Striate Cortex 1 R | 0.000 | DEF C: Parahippocampal Cortex 1 R | 0.2 |
| DA A: TEMP Occipital 1 R | 0.000 | LIM A: Temp Pole 2 L | 0.2 |
| DEF C: Parahippocampal Cortex 1 R | 0.000 | DEF C: Parahippocampal Cortex 1 L | 0.2 |
| TP 3 R | 0.000 | TP 1 L | 0.1 |

**Table S4. Dynamic metabolic State 3.** Rank order of regional clustering and centrality for State 3. Blue shading is top 25% of regions.

| State 3: Clustering | | State 3: Centrality | |
|---|---|---|---|
| DA B: Post Cent 2 R | 0.322 | CON B: Lateral Prefrontal Cortexv 1 R | 31.4 |
| DEF B: Ventral Prefrontal Cortex 1 L | 0.320 | DEF B: Ventral Prefrontal Cortex 2 L | 27.1 |
| SOM B: S2 2 R | 0.316 | CON B: Lateral Prefrontal Cortexv 1 L | 26.9 |
| SVA A: Insula: 2 L | 0.307 | DEF B: Dorsal Prefrontal Cortex 1 L | 25.3 |
| SVA B: Medial Posterior Prefrontal 1 L | 0.306 | DEF B: Inferior Parietal Lobule 1 L | 24.6 |
| SOM B: Cent 1 L | 0.302 | DEF B: Dorsal Prefrontal Cortex 1 R | 23.8 |
| DA A: Parietal Occipital 1 L | 0.294 | CON A: Lateral Prefrontal Cortex 1 R | 23.3 |
| TP 1 R | 0.292 | CON A: Lateral Prefrontal Cortex 1 L | 22.1 |
| SVA A: Parietal Operculum 1 L | 0.291 | VIS Cent: Extra Striate Cortex 2 R | 22.0 |
| DA B: Frontal Eye Fields 1 R | 0.288 | DEF B: Ventral Prefrontal Cortex 2 R | 21.5 |
| SVA A: Insula: 1 R | 0.287 | SVA B: Lateral Prefrontal Cortex 1 L | 20.6 |
| DEF B: Temp 1 L | 0.284 | CON B: inferior parietal lobule 1 R | 19.6 |
| VIS Perhiph: Striate Cortex Calcarine 1 R | 0.283 | CON B: Lateral Prefrontal Cortexd 1 R | 19.5 |
| DEF B: Ventral Prefrontal Cortex 1 R | 0.282 | VIS Cent: Extra Striate Cortex 2 L | 18.7 |
| DA A: TEMP Occipital 1 R | 0.279 | SVA B: Lateral Prefrontal Cortex 1 R | 18.7 |
| LIM A Temp Pole 1 L | 0.273 | DA A: Superior Parietal Lobule 1 L | 18.6 |
| SVA A: Parietal Operculum 1 R | 0.273 | SOM B: Auditory 1 L | 18.1 |
| SVA B: Medial Posterior Prefrontal 1 R | 0.272 | CON B: Temp 1 R | 17.6 |
| TP 1 L | 0.270 | SOM B: Auditory 1 R | 17.6 |
| DA B: Frontal Eye Fields 1 L | 0.267 | CON A: Lateral Prefrontal Cortex 2 R | 16.7 |
| CON C: Precuneus 1 L | 0.267 | CON A: Intraparietal Sulcus 1 L | 16.5 |
| DA B: Post Cent 1 R | 0.267 | VIS Cent: Extra Striate Cortex 3 L | 16.4 |
| SVA B: Inferior Parietal Lobule 1 R | 0.266 | LIM B: Orbital Frontal Cortex 0 R | 15.1 |
| DEF A: Medial Prefrontal Cortex 1 L | 0.265 | VIS Perhiph: Extra Striate Superior 1 R | 14.8 |
| DEF A: Precuneus Posterior Cingulate Cortex 1 R | 0.263 | DA A: Parietal Occipital 1 R | 14.1 |
| DA B: Post Cent 1 L | 0.261 | VIS Cent: Extra Striate Cortex 3 R | 13.9 |
| DA B: Post Cent 3 L | 0.260 | LIM B: Orbital Frontal Cortex 1 L | 13.1 |
| DEF B: Lateral Prefrontal Cortex 1 L | 0.260 | DEF A: Medial Prefrontal Cortex 1 R | 12.8 |
| DEF B: Temp 2 L | 0.258 | VIS Cent: Striate Cortex 1 L | 12.5 |
| SVA A: Parietal Medial 1 L | 0.258 | SOM A: 2 L | 11.2 |
| SVA A: Frontal Medial 1 R | 0.257 | DEF A: Inferior Parietal Lobule 1 R | 11.1 |
| CON A: Lateral Prefrontal Cortex 2 L | 0.257 | CON C: Precuneus 1 R | 10.3 |
| DA A: Superior Parietal Lobule 1 R | 0.257 | SVA B: Inferior Parietal Lobule 1 R | 10.2 |
| VIS Cent: Extra Striate Cortex 1 L | 0.256 | DA A: TEMP Occipital 1 L | 10.1 |
| SOM B: S2 2 L | 0.256 | DEF A: Medial Prefrontal Cortex 1 L | 10.1 |
| SOM A: 1 L | 0.254 | VIS Perhiph: Extra Striate CortexSup 1 L | 10.1 |
| CON A: Intraparietal Sulcus 1 R | 0.254 | LIM A Temp Pole 1 L | 10.0 |
| CON C: Cingulate Posterior 1 L | 0.254 | DEF B: Temp 1 L | 10.0 |
| DEF A: Inferior Parietal Lobule 1 R | 0.252 | DA A: Superior Parietal Lobule 1 R | 9.9 |
| DA A: TEMP Occipital 1 L | 0.252 | CON A: Lateral Prefrontal Cortex 2 L | 9.9 |
| CON C: Cingulate Posterior 1 R | 0.252 | DEF A: Dorsal Prefrontal Cortex 1 L | 9.9 |
| SVA A: Frontal Medial 1 L | 0.248 | DA A: TEMP Occipital 1 R | 9.8 |
| VIS Cent: Extra Striate Cortex 3 R | 0.247 | SOM B: S2 2 L | 9.7 |
| VIS Cent: Extra Striate Cortex 1 R | 0.247 | DEF A: Dorsal Prefrontal Cortex 1 R | 9.7 |
| DEF A: Dorsal Prefrontal Cortex 1 L | 0.247 | DEF B: Lateral Prefrontal Cortex 1 L | 9.6 |
| VIS Cent: Striate Cortex 1 L | 0.246 | LIM A: Temp Pole 1 R | 9.3 |
| SOM B: Cent 1 R | 0.246 | SOM A: 1 L | 9.0 |
| LIM B: Orbital Frontal Cortex 1 L | 0.242 | DEF B: Ventral Prefrontal Cortex 1 R | 8.3 |
| SVA A: Parietal Medial 1 R | 0.242 | SVA A: Parietal Operculum 1 R | 8.2 |
| VIS Perhiph: Extra Striate CortexSup 1 L | 0.242 | SVA A: Frontal Medial 1 L | 7.9 |
| SOM A: 4 R | 0.241 | SVA A: Parietal Operculum 1 L | 7.7 |
| DA A: Parietal Occipital 1 R | 0.241 | DA A: Parietal Occipital 1 L | 7.7 |
| DEF A: Dorsal Prefrontal Cortex 1 R | 0.240 | SOM B: Cent 1 R | 7.5 |
| LIM A: Temp Pole 1 R | 0.238 | VIS Cent: Extra Striate Cortex 1 L | 7.5 |
| LIM B: Orbital Frontal Cortex 0 R | 0.236 | SOM B: Cent 1 L | 7.4 |
| VIS Perhiph: Extra Striate Superior 1 R | 0.235 | DA B: Frontal Eye Fields 1 L | 7.2 |
| DEF A: Precuneus Posterior Cingulate Cortex1 L | 0.234 | DA B: Post Cent 3 L | 7.2 |
| DEF A: Medial Prefrontal Cortex 1 R | 0.233 | TP 1 L | 6.5 |
| CON C: Precuneus 1 R | 0.231 | DEF A: Precuneus Posterior Cingulate Cortex1 L | 6.3 |
| VIS Cent: Extra Striate Cortex 3 L | 0.225 | DEF B: Temp 2 L | 5.8 |
| VIS Perhiph: Extra Striate Inferior 1 R | 0.221 | DA B: Frontal Eye Fields 1 R | 5.8 |
| VIS Perhiph: Striate Cortex Calcarine 1 L | 0.216 | CON C: Precuneus 1 L | 5.8 |
| VIS Cent: Extra Striate Cortex 2 L | 0.209 | CON A: Intraparietal Sulcus 1 R | 5.1 |
| TP 2 R | 0.207 | VIS Cent: Extra Striate Cortex 1 R | 5.1 |
| CON A: Lateral Prefrontal Cortex 2 R | 0.205 | DEF A: Precuneus Posterior Cingulate Cortex 1 R | 4.7 |
| SOM A: 2 R | 0.204 | VIS Perhiph: Striate Cortex Calcarine 1 R | 4.4 |
| CON B: Temp 1 R | 0.204 | SVA A: Frontal Medial 1 R | 4.2 |
| SOM A: 2 L | 0.203 | DA B: Post Cent 1 L | 4.0 |
| SOM B: Auditory 1 R | 0.201 | SOM B: S2 2 R | 4.0 |
| DEF B: Ventral Prefrontal Cortex 2 R | 0.199 | SVA A: Insula: 1 R | 4.0 |
| VIS Cent: Extra Striate Cortex 2 R | 0.197 | SVA A: Parietal Medial 1 R | 3.8 |
| DA A: Superior Parietal Lobule 1 L | 0.195 | VIS Perhiph: Striate Cortex Calcarine 1 L | 3.8 |
| CON B: inferior parietal lobule 1 R | 0.194 | SOM A: 4 R | 3.7 |
| SVA B: Lateral Prefrontal Cortex 1 R | 0.193 | DA B: Post Cent 1 R | 3.5 |
| CON A: Intraparietal Sulcus 1 L | 0.193 | DEF B: Ventral Prefrontal Cortex 1 L | 3.3 |
| SOM B: Auditory 1 L | 0.191 | SVA B: Medial Posterior Prefrontal 1 L | 3.0 |
| SVA B: Lateral Prefrontal Cortex 1 L | 0.188 | VIS Perhiph: Extra Striate Inferior 1 R | 2.6 |
| CON A: Lateral Prefrontal Cortex 1 L | 0.183 | TP 2 R | 2.2 |
| CON B: Lateral Prefrontal Cortexd 1 R | 0.183 | SVA B: Medial Posterior Prefrontal 1 R | 2.0 |
| SOM A: 3 R | 0.183 | SOM A: 2 R | 2.0 |
| DEF B: Inferior Parietal Lobule 1 L | 0.177 | SVA A: Parietal Medial 1 L | 1.9 |
| CON A: Lateral Prefrontal Cortex 1 R | 0.172 | CON C: Cingulate Posterior 1 R | 1.9 |
| DEF B: Ventral Prefrontal Cortex 2 L | 0.170 | DA B: Post Cent 2 R | 1.7 |
| CON B: Lateral Prefrontal Cortexv 1 L | 0.169 | CON C: Cingulate Posterior 1 L | 1.7 |
| DEF B: Dorsal Prefrontal Cortex 1 R | 0.168 | TP 1 R | 1.6 |
| DEF B: Dorsal Prefrontal Cortex 1 L | 0.160 | SVA A: Insula: 2 L | 1.3 |
| CON B: Lateral Prefrontal Cortexv 1 R | 0.155 | TP 3 R | 1.1 |
| TP 3 R | 0.102 | SOM A: 3 R | 0.8 |
| VIS Perhiph: Extra Striate Inferior 1 L | 0.000 | SOM A: 1 R | 0.6 |
| SOM B: S2 1 L | 0.000 | VIS Perhiph: Extra Striate Inferior 1 L | 0.6 |
| DA B: Post Cent 2 L | 0.000 | DA B: Post Cent 2 L | 0.6 |
| SVA A: Insula: 1 L | 0.000 | DEF C: RetroSuperior Parietal Lobuleenial 1 R | 0.6 |
| LIM A: Temp Pole 2 L | 0.000 | DEF C: Parahippocampal Cortex 1 R | 0.5 |
| CON C: Precuneus 2 L | 0.000 | CON C: Precuneus 2 L | 0.3 |
| DEF C: RetroSuperior Parietal Lobuleenial 1 L | 0.000 | DEF C: RetroSuperior Parietal Lobuleenial 1 L | 0.3 |
| DEF C: Parahippocampal Cortex 1 L | 0.000 | LIM A: Temp Pole 2 L | 0.3 |
| SOM A: 1 R | 0.000 | SOM B: S2 1 L | 0.3 |
| SOM B: S2 1 R | 0.000 | SVA A: Insula: 1 L | 0.0 |
| DEF C: RetroSuperior Parietal Lobuleenial 1 R | 0.000 | DEF C: Parahippocampal Cortex 1 L | 0.0 |
| DEF C: Parahippocampal Cortex 1 R | 0.000 | SOM B: S2 1 R | 0.0 |

**Table S5. Dynamic metabolic State 4.** Rank order of regional clustering and centrality for State 4. Blue shading is top 25% of regions.

| State 4: Clustering | | State 4: Centrality | |
|---|---|---|---|
| DEF C: Parahippocampal Cortex 1 L | 0.024 | DEF B: Ventral Prefrontal Cortex 2 L | 2.1 |
| VIS Perhiph: Extra Striate Inferior 1 L | 0.022 | DEF B: Temp 1 L | 2.1 |
| DA A: TEMP Occipital 1 R | 0.021 | DA A: TEMP Occipital 1 L | 1.9 |
| VIS Cent: Extra Striate Cortex 1 R | 0.021 | VIS Cent: Extra Striate Cortex 2 R | 1.9 |
| VIS Cent: Extra Striate Cortex 2 L | 0.021 | VIS Cent: Extra Striate Cortex 3 L | 1.8 |
| DEF C: RetroSuperior Parietal Lobuleenial 1 L | 0.021 | DEF A: Precuneus Posterior Cingulate Cortex1 L | 1.7 |
| DEF C: Parahippocampal Cortex 1 R | 0.020 | CON B: Lateral Prefrontal Cortexv 1 R | 1.6 |
| DEF A: Dorsal Prefrontal Cortex 1 L | 0.020 | DEF B: Inferior Parietal Lobule 1 L | 1.6 |
| VIS Perhiph: Extra Striate Inferior 1 R | 0.020 | CON B: Lateral Prefrontal Cortexv 1 L | 1.6 |
| LIM B: Orbital Frontal Cortex 1 L | 0.020 | LIM A Temp Pole 1 L | 1.6 |
| LIM A: Temp Pole 1 R | 0.020 | CON B: Temp 1 R | 1.6 |
| LIM A: Temp Pole 2 L | 0.020 | LIM B: Orbital Frontal Cortex 0 R | 1.6 |
| DEF B: Temp 2 L | 0.019 | SVA B: Lateral Prefrontal Cortex 1 L | 1.5 |
| VIS Cent: Extra Striate Cortex 1 L | 0.019 | VIS Perhiph: Striate Cortex Calcarine 1 R | 1.5 |
| TP 1 R | 0.019 | VIS Cent: Extra Striate Cortex 3 R | 1.5 |
| DEF C: RetroSuperior Parietal Lobuleenial 1 R | 0.018 | LIM B: Orbital Frontal Cortex 1 L | 1.5 |
| SVA A: Frontal Medial 1 L | 0.017 | DA A: Parietal Occipital 1 R | 1.5 |
| VIS Cent: Extra Striate Cortex 2 R | 0.017 | VIS Perhiph: Striate Cortex Calcarine 1 L | 1.5 |
| DEF B: Lateral Prefrontal Cortex 1 L | 0.017 | VIS Perhiph: Extra Striate Inferior 1 R | 1.5 |
| VIS Perhiph: Striate Cortex Calcarine 1 R | 0.017 | VIS Cent: Extra Striate Cortex 1 R | 1.5 |
| VIS Perhiph: Extra Striate CortexSup 1 L | 0.017 | CON A: Intraparietal Sulcus 1 L | 1.5 |
| DA A: TEMP Occipital 1 L | 0.016 | DA A: Superior Parietal Lobule 1 L | 1.5 |
| DEF A: Precuneus Posterior Cingulate Cortex 1 R | 0.016 | CON C: Precuneus 1 R | 1.5 |
| DEF B: Ventral Prefrontal Cortex 1 L | 0.016 | DEF B: Dorsal Prefrontal Cortex 1 L | 1.5 |
| SVA A: Parietal Operculum 1 L | 0.016 | VIS Cent: Extra Striate Cortex 2 L | 1.4 |
| SOM B: S2 2 L | 0.016 | CON C: Precuneus 2 L | 1.4 |
| SVA A: Parietal Medial 1 L | 0.016 | DA A: TEMP Occipital 1 R | 1.4 |
| SVA B: Medial Posterior Prefrontal 1 R | 0.016 | VIS Perhiph: Extra Striate Inferior 1 L | 1.4 |
| VIS Perhiph: Striate Cortex Calcarine 1 L | 0.016 | CON A: Lateral Prefrontal Cortex 1 L | 1.4 |
| DA B: Post Cent 1 R | 0.016 | CON A: Lateral Prefrontal Cortex 1 R | 1.4 |
| SVA A: Insula: 1 L | 0.016 | CON B: Lateral Prefrontal Cortexd 1 R | 1.4 |
| LIM B: Orbital Frontal Cortex 0 R | 0.015 | SOM A: 1 L | 1.4 |
| CON C: Precuneus 1 L | 0.015 | VIS Cent: Striate Cortex 1 L | 1.4 |
| SOM B: Auditory 1 L | 0.015 | SOM B: Auditory 1 R | 1.3 |
| DA A: Parietal Occipital 1 L | 0.015 | SOM A: 2 L | 1.3 |
| VIS Cent: Extra Striate Cortex 3 R | 0.015 | CON C: Precuneus 1 L | 1.3 |
| VIS Cent: Extra Striate Cortex 3 L | 0.015 | VIS Perhiph: Extra Striate Superior 1 R | 1.3 |
| CON B: Temp 1 R | 0.015 | SOM A: 4 R | 1.3 |
| DA B: Frontal Eye Fields 1 R | 0.015 | VIS Perhiph: Extra Striate CortexSup 1 L | 1.3 |
| DA B: Post Cent 2 L | 0.015 | TP 1 L | 1.3 |
| CON A: Lateral Prefrontal Cortex 2 L | 0.015 | SVA A: Parietal Operculum 1 R | 1.3 |
| SVA B: Medial Posterior Prefrontal 1 L | 0.015 | TP 3 R | 1.3 |
| SVA A: Insula: 2 L | 0.015 | SVA B: Lateral Prefrontal Cortex 1 R | 1.3 |
| TP 1 L | 0.015 | LIM A: Temp Pole 1 R | 1.3 |
| VIS Perhiph: Extra Striate Superior 1 R | 0.015 | DEF C: Parahippocampal Cortex 1 R | 1.2 |
| VIS Cent: Striate Cortex 1 L | 0.015 | VIS Cent: Extra Striate Cortex 1 L | 1.2 |
| CON B: Lateral Prefrontal Cortexv 1 L | 0.015 | DA A: Superior Parietal Lobule 1 R | 1.2 |
| DEF B: Temp 1 L | 0.015 | TP 2 R | 1.2 |
| SVA A: Parietal Medial 1 R | 0.014 | SOM B: S2 2 L | 1.2 |
| DEF A: Precuneus Posterior Cingulate Cortex1 L | 0.014 | SVA A: Insula: 2 L | 1.2 |
| SVA B: Inferior Parietal Lobule 1 R | 0.014 | DEF B: Lateral Prefrontal Cortex 1 L | 1.2 |
| CON A: Lateral Prefrontal Cortex 1 L | 0.014 | DEF A: Inferior Parietal Lobule 1 R | 1.2 |
| SOM B: Auditory 1 R | 0.014 | DEF B: Ventral Prefrontal Cortex 1 R | 1.2 |
| LIM A Temp Pole 1 L | 0.014 | DEF B: Dorsal Prefrontal Cortex 1 R | 1.2 |
| CON A: Intraparietal Sulcus 1 L | 0.014 | DEF B: Ventral Prefrontal Cortex 2 R | 1.2 |
| DEF B: Ventral Prefrontal Cortex 1 R | 0.014 | SOM B: Auditory 1 L | 1.2 |
| DEF B: Dorsal Prefrontal Cortex 1 R | 0.014 | CON A: Intraparietal Sulcus 1 R | 1.2 |
| SOM A: 2 L | 0.014 | DA A: Parietal Occipital 1 L | 1.2 |
| SOM B: S2 1 R | 0.014 | DEF A: Dorsal Prefrontal Cortex 1 R | 1.1 |
| TP 2 R | 0.014 | DA B: Post Cent 1 L | 1.1 |
| CON A: Intraparietal Sulcus 1 R | 0.014 | DA B: Frontal Eye Fields 1 L | 1.1 |
| DA B: Frontal Eye Fields 1 L | 0.014 | CON A: Lateral Prefrontal Cortex 2 R | 1.1 |
| CON B: Lateral Prefrontal Cortexv 1 R | 0.014 | LIM A: Temp Pole 2 L | 1.1 |
| CON C: Precuneus 1 R | 0.014 | CON C: Cingulate Posterior 1 L | 1.1 |
| DEF A: Dorsal Prefrontal Cortex 1 R | 0.014 | DEF C: RetroSuperior Parietal Lobuleenial 1 R | 1.1 |
| DA B: Post Cent 2 R | 0.014 | DEF C: RetroSuperior Parietal Lobuleenial 1 L | 1.0 |
| CON A: Lateral Prefrontal Cortex 1 R | 0.014 | CON A: Lateral Prefrontal Cortex 2 L | 1.0 |
| DA B: Post Cent 1 L | 0.014 | SOM A: 2 R | 1.0 |
| DEF A: Inferior Parietal Lobule 1 R | 0.013 | DEF A: Precuneus Posterior Cingulate Cortex 1 R | 1.0 |
| SOM B: S2 1 L | 0.013 | DEF B: Ventral Prefrontal Cortex 1 L | 1.0 |
| TP 3 R | 0.013 | DA B: Post Cent 2 R | 1.0 |
| SOM A: 4 R | 0.013 | SVA A: Parietal Medial 1 L | 1.0 |
| DA B: Post Cent 3 L | 0.013 | DA B: Post Cent 3 L | 1.0 |
| SOM B: S2 2 R | 0.013 | SVA B: Medial Posterior Prefrontal 1 L | 1.0 |
| CON C: Cingulate Posterior 1 L | 0.013 | SVA A: Parietal Medial 1 R | 1.0 |
| DEF B: Ventral Prefrontal Cortex 2 L | 0.013 | SVA B: Inferior Parietal Lobule 1 R | 1.0 |
| DEF B: Dorsal Prefrontal Cortex 1 L | 0.013 | SVA A: Insula: 1 R | 1.0 |
| SOM A: 1 R | 0.013 | CON C: Cingulate Posterior 1 R | 1.0 |
| SVA B: Lateral Prefrontal Cortex 1 L | 0.013 | TP 1 R | 1.0 |
| CON C: Precuneus 2 L | 0.013 | SVA A: Parietal Operculum 1 L | 1.0 |
| DEF B: Ventral Prefrontal Cortex 2 R | 0.013 | DEF A: Medial Prefrontal Cortex 1 R | 0.9 |
| DEF B: Inferior Parietal Lobule 1 L | 0.013 | SVA A: Frontal Medial 1 R | 0.9 |
| DA A: Superior Parietal Lobule 1 L | 0.013 | DA B: Post Cent 2 L | 0.9 |
| DEF A: Medial Prefrontal Cortex 1 L | 0.013 | DEF A: Medial Prefrontal Cortex 1 L | 0.9 |
| SVA A: Insula: 1 R | 0.012 | SVA A: Frontal Medial 1 L | 0.9 |
| CON C: Cingulate Posterior 1 R | 0.012 | DEF A: Dorsal Prefrontal Cortex 1 L | 0.9 |
| DA A: Parietal Occipital 1 R | 0.012 | SVA A: Insula: 1 L | 0.8 |
| SVA A: Frontal Medial 1 R | 0.012 | CON B: inferior parietal lobule 1 R | 0.8 |
| CON B: inferior parietal lobule 1 R | 0.012 | SOM A: 1 R | 0.8 |
| SVA B: Lateral Prefrontal Cortex 1 R | 0.012 | DEF C: Parahippocampal Cortex 1 L | 0.8 |
| DEF A: Medial Prefrontal Cortex 1 R | 0.012 | DA B: Post Cent 1 R | 0.8 |
| SOM A: 1 L | 0.012 | SOM B: Cent 1 L | 0.8 |
| SOM B: Cent 1 L | 0.012 | DA B: Frontal Eye Fields 1 R | 0.8 |
| SVA A: Parietal Operculum 1 R | 0.012 | SOM B: Cent 1 R | 0.8 |
| SOM A: 2 R | 0.011 | SOM B: S2 1 L | 0.8 |
| DA A: Superior Parietal Lobule 1 R | 0.011 | SOM B: S2 2 R | 0.8 |
| SOM A: 3 R | 0.011 | SOM A: 3 R | 0.8 |
| SOM B: Cent 1 R | 0.011 | SVA B: Medial Posterior Prefrontal 1 R | 0.7 |
| CON A: Lateral Prefrontal Cortex 2 R | 0.011 | DEF B: Temp 2 L | 0.7 |
| CON B: Lateral Prefrontal Cortexd 1 R | 0.011 | SOM B: S2 1 R | 0.7 |

##### *2.4 Timeseries Standard Deviation and Sample Entropy*

While both standard deviation and entropy of neuroimaging signals describe aspects of brain variability, they capture different properties (Figure S2). The standard deviation reflects the magnitude of fluctuations - a global, summary measure of how much the signal deviates from the mean over time. For example, in fPET, a high standard deviation reflects greater rises and falls in glucose uptake. However, standard deviation does not capture the temporal structure of the fluctuations. Entropy, by contrast, measures the complexity or unpredictability of the signal. A high-entropy fPET signal reflects a more irregular, complex pattern of metabolic activity, indicative of rich and dynamic glucose use. Two signals may have the same standard deviation but differ in entropy - or vice versa - depending on the structure and shape of the timeseries. Therefore, standard deviation and entropy provide complementary measures of the brain's capacity for dynamic state transitions and information transfer.

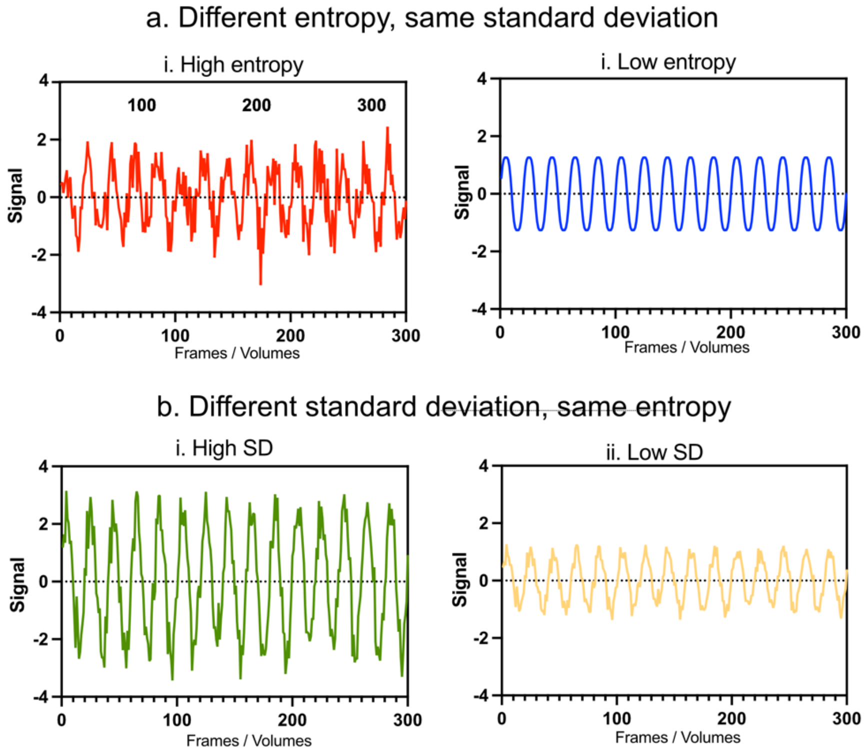


**Fig S2. Simulation of entropy and standard deviation of neuroimaging timeseries.** Signals with different entropy but same standard deviation of the timeseries (a) and signals with different standard deviation but the same entropy of the timeseries (b).

##### *2.5 Neurochemical system alignment with metabolic network states*

**Table S6. Neurochemical system alignment with metabolic network states.** Correlations (r) between centrality and neurochemical system maps from *neuromaps* (A), and loadings from principal component analysis (B).

| | Centrality Map Correlation (r) | | | | Transmitter-to-PC Scores | | |
|---|---|---|---|---|---|---|---|
| **System Family / Biological Target** | **State 1** | **State 2** | **State 3** | **State 4** | **PC1** | **PC2** | **PC3** |
| **Serotonin (5-HT) System** | | | | | | | |
| Serotonin 5-HT1A receptor (Savli, 2012) | 0.35 | 0.02 | 0.21 | 0.46 | 0.70 | -0.90 | 0.38 |
| Serotonin 5-HT2A receptor (Savli, 2012) | 0.54 | 0.16 | 0.44 | 0.70 | 0.77 | -0.89 | 0.34 |
| Serotonin 5-HT6 receptor GSK215083 (Radnakrishnan, 2018) | 0.47 | 0.23 | 0.44 | 0.65 | 0.78 | -0.80 | 0.42 |
| Serotonin 5-HT1B receptor P943 (Gallezot, 2010) | 0.36 | 0.34 | 0.46 | 0.29 | 0.36 | 0.08 | -0.92 |
| Serotonin transported (SERT) 5HTT (Savli, 2012) | -0.07 | -0.33 | -0.29 | -0.24 | 0.23 | -0.86 | -0.16 |
| Serotonin transported (SERT) MADAM (Fazio, 2016) | 0.10 | -0.10 | -0.02 | 0.03 | 0.51 | -0.98 | -0.04 |
| **Dopamine (DA) System** | | | | | | | |
| Dopamine $D_1$ receptor D1 (Kaller, 2017) | 0.39 | 0.04 | 0.16 | 0.46 | 0.55 | -0.90 | 0.43 |
| Dopamine $D_2$ receptor D2 (Alarkurtti, 2015) | 0.02 | 0.15 | 0.10 | 0.13 | -0.31 | 0.78 | 0.51 |
| Extrastriatal $D_2/_3$ receptor FLB457 (Sandiego, 2015) | -0.11 | -0.50 | -0.27 | -0.51 | 0.31 | -0.60 | -0.76 |
| Extrastriatal $D_2/_3$ receptor FALLYPRIDE (Jaworska, 2020) | -0.24 | -0.47 | -0.36 | -0.63 | 0.00 | -0.27 | -0.88 |
| Dopamine transported (DAT) DAT (Aghourian, 2017) | 0.00 | 0.31 | 0.05 | 0.22 | -0.62 | 0.73 | 0.68 |
| Dopamine transported (DAT) FEPE2I (Sasaki, 2012) | -0.03 | 0.00 | -0.04 | 0.00 | -0.57 | 0.30 | 0.90 |
| Mixed Dopamine/Serotonin transported FPCIT (Dukart, 2018) | -0.26 | -0.51 | -0.44 | -0.67 | -0.08 | -0.31 | -0.79 |
| **Norepinephrine (NE) System** | | | | | | | |
| Norepinephrine transported (NET) MRB (Ding, 2010) | -0.14 | 0.34 | 0.09 | -0.02 | -0.64 | 1.00 | 0.03 |
| Norepinephrine transported (NET) METHYLREBOXETINE (Hesse, 2017) | 0.05 | 0.20 | 0.08 | 0.35 | -0.04 | 0.13 | 0.96 |
| **GABAergic System** | | | | | | | |
| $GABA_A$ receptor GABAA (Dukart, 2018) | 0.51 | 0.14 | 0.38 | 0.73 | 0.70 | -0.85 | 0.45 |
| $GABA_A$ Benzodiazepine Site RO154513 (Lukow, 2022) | 0.37 | 0.16 | 0.25 | 0.52 | 0.57 | -0.79 | 0.58 |
| $GABA_A$ receptor Availability PS13 (Kim, 2020) | 0.13 | -0.35 | -0.02 | 0.02 | 0.76 | -0.98 | -0.14 |
| **Opioid System** | | | | | | | |
| μ-Opioid receptor CARFENTANIL (Kantonen, 2020) | 0.03 | 0.20 | 0.08 | -0.16 | -0.65 | 0.74 | -0.59 |
| κ-Opioid receptor LY2795050 (Vijay, 2018) | 0.09 | 0.35 | 0.23 | 0.12 | -0.61 | 0.99 | -0.13 |
| **Cholinergic System** | | | | | | | |
| $M_1$ Muscarinic ACh receptor LSN3172176 (Naganawa, 2020) | 0.44 | 0.19 | 0.39 | 0.63 | 0.75 | -0.80 | 0.46 |
| $\alpha_4\beta_2$ Nicotinic ACh receptor FLUBATINE (Hillmer, 2016) | -0.21 | 0.05 | -0.06 | -0.55 | -0.53 | 0.69 | -0.69 |
| **Glutamatergic / Excitation Systems** | | | | | | | |
| $mGluR_5$ Metabotropic Glutamate mGluR5 (Smart, 2019) | 0.36 | 0.28 | 0.31 | 0.58 | 0.47 | -0.56 | 0.78 |
| NMDA receptor Ion Channel Site GE179 (Galovic, 2021) | 0.39 | 0.26 | 0.33 | 0.64 | 0.53 | -0.62 | 0.72 |
| **Baselines & Microglial Targets** | | | | | | | |
| Cannabinoid $CB_1$ receptor CB1 (Laurikainen, 2018) | 0.37 | 0.48 | 0.50 | 0.53 | 0.23 | 0.42 | 0.48 |
| Cannabinoid $CB_1$ receptor OMAR (Normandin, 2015) | 0.34 | 0.21 | 0.35 | 0.36 | 0.94 | -0.87 | -0.03 |
| Histamine $H_3$ receptor GSK189254 (Gallezot, 2017) | -0.01 | 0.24 | 0.06 | -0.11 | -0.76 | 0.89 | -0.34 |
| Histone Deacetylase (HDAC) MARTINOSTAT (Wey, 2016) | 0.46 | 0.35 | 0.48 | 0.62 | 0.78 | -0.67 | 0.49 |
| Mean Cerebral Blood Flow MEANCBF (Satterthwaite, 2014) | 0.40 | 0.34 | 0.41 | 0.48 | 0.79 | -0.68 | 0.49 |
| Synaptic Vesicle Glycoprotein 2A UCBJ (Finnema, 2016) | 0.38 | 0.14 | 0.30 | 0.56 | 0.68 | -0.79 | 0.52 |
| TSPO / Microglial Activation PBR28 (Lois, 2018) | -0.06 | -0.38 | -0.24 | -0.40 | 0.21 | -0.61 | -0.69 |

Note: Shaded cells indicate statistically significant state-receptor associations (p-FDR < 0.05) based on permutation testing (n=1,000). The large number of cortical vertices (~20,000) provides high statistical power to detect even subtle but consistent spatial alignments between metabolic network states and receptor distributions.

##### *2.6 Region Glucodynamics*

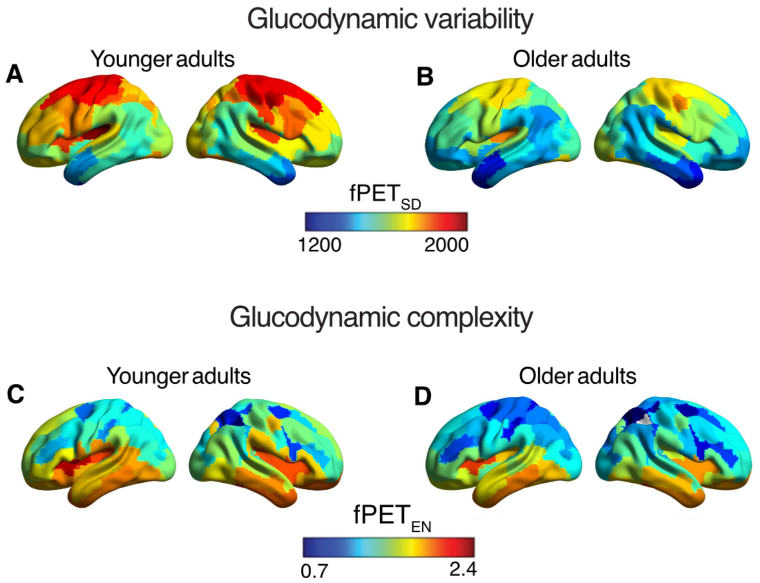


**Figure S3. Age group differences in dynamic metabolic regional brain signal measures.** Younger and older adult mean regional $fPET_{SD}$ (A) and $fPET_{ENT}$ (B).

The dynamic regional brain signal measures for the younger and older adults are shown in Fig S3. Older adults had reduced $fPET_{SD}$ in 86 regions and reduced $fPET_{EN}$ in 71 regions. There were no regions in which older adults had greater glucodynamic variability or complexity than younger adults (the GLMs are available in the Supp Tables S7 and S8). The effect sizes of the age group differences revealed distinct spatial topographies for $fPET_{EN}$ and $fPET_{SD}$. The largest age group differences in glucodynamic complexity were distributed across primary sensory and higher-order cognitive networks, showing their largest differences in somatomotor and visual systems as well as bilateral dorsal and ventral prefrontal regions, right medial prefrontal cortex and left temporal cortex. This was accompanied by lower complexity among older adults in key nodes of the salience ventral attention (bilateral insula and parietal operculum) and control (bilateral precuneus and right inferior parietal lobule) networks.

Conversely, age-related differences in $fPET_{SD}$ exhibited a pattern of lower variability among older adults in regions of the control network, including bilateral ventrolateral and dorsolateral prefrontal regions. Older adults also had lower variability in regions of the default network, including bilateral medial, ventral and dorsal prefrontal cortices, as well as bilateral posterior cingulate and left temporal structures. Lower glucodynamic variability in older adults extended into paralimbic and limbic regions, including the bilateral temporal poles and right orbitofrontal cortex, as well as frontoparietal regions of the salience system.

These results suggest that ageing shapes glucodynamic brain signal features, dampening the dynamic range, complexity and information processing capacity of core functional networks.

**Table S7. Whole sample and younger and older adult group regional entropy.** Mean and standard deviation of 100 region $fPET_{EN}$ values and GLMs of age group differences.

| | Younger | | Older | | Younger vs older | | | | Head motion | | | |
|---|---|---|---|---|---|---|---|---|---|---|---|---|
| | Mean | SD | Mean | SD | F | p | p-FDR | η2p | F | p | p-FDR | η2p |
| Visual Central: Extra Striate Cortex 1 L | 2.12 | 0.10 | 2.10 | 0.14 | 0.2 | 0.693 | 0.721 | 0.002 | 0.26 | 0.613 | 0.947 | 0.003 |
| Visual Central: Extra Striate Cortex 2 L | 2.06 | 0.07 | 1.97 | 0.09 | 29.7 | 0.000 | 0.000 | 0.266 | 0.02 | 0.898 | 0.989 | 0.000 |
| Visual Central: Striate Cortex 1 L | 1.99 | 0.08 | 1.91 | 0.10 | 15.6 | 0.000 | 0.000 | 0.160 | 0.22 | 0.638 | 0.947 | 0.003 |
| Visual Central: Extra Striate Cortex 3 L | 1.97 | 0.08 | 1.90 | 0.08 | 12.3 | 0.001 | 0.001 | 0.130 | 0.02 | 0.886 | 0.989 | 0.000 |
| Visual Peripheral: Extra Striate Inferior 1 L | 2.23 | 0.06 | 2.17 | 0.11 | 11.3 | 0.001 | 0.002 | 0.121 | 0.54 | 0.466 | 0.847 | 0.007 |
| Visual Peripheral: Striate Cortex Calcarine 1 L | 1.99 | 0.12 | 1.92 | 0.11 | 7.1 | 0.009 | 0.012 | 0.080 | 0.00 | 0.949 | 0.989 | 0.000 |
| Visual Peripheral: Extra Striate CortexSup 1 L | 1.99 | 0.08 | 1.89 | 0.10 | 23.0 | 0.000 | 0.000 | 0.219 | 0.01 | 0.934 | 0.989 | 0.000 |
| Somatomotor A: 1 L | 1.98 | 0.07 | 1.86 | 0.13 | 22.1 | 0.000 | 0.000 | 0.212 | 6.07 | 0.016 | 0.469 | 0.069 |
| Somatomotor A: 2 L | 1.97 | 0.07 | 1.87 | 0.11 | 19.8 | 0.000 | 0.000 | 0.194 | 1.16 | 0.285 | 0.712 | 0.014 |
| Somatomotor B: Auditory 1 L | 2.04 | 0.07 | 1.95 | 0.08 | 26.5 | 0.000 | 0.000 | 0.244 | 0.02 | 0.901 | 0.989 | 0.000 |
| Somatomotor B: S2 1 L | 2.27 | 0.06 | 2.17 | 0.08 | 42.1 | 0.000 | 0.000 | 0.339 | 1.17 | 0.282 | 0.712 | 0.014 |
| Somatomotor B: S2 2 L | 2.21 | 0.05 | 1.99 | 0.10 | 130.6 | 0.000 | 0.000 | 0.614 | 0.83 | 0.366 | 0.796 | 0.010 |
| Somatomotor B: Central 1 L | 2.01 | 0.09 | 1.96 | 0.10 | 4.1 | 0.047 | 0.053 | 0.047 | 1.66 | 0.201 | 0.682 | 0.020 |
| Dorsal Attention A: Temporal Occipital 1 L | 2.21 | 0.06 | 2.18 | 0.07 | 4.9 | 0.029 | 0.034 | 0.057 | 0.71 | 0.401 | 0.809 | 0.009 |
| Dorsal Attention A: Parietal Occipital 1 L | 2.00 | 0.13 | 1.98 | 0.10 | 0.9 | 0.338 | 0.363 | 0.011 | 0.45 | 0.504 | 0.890 | 0.005 |
| Dorsal Attention A: Superior Parietal Lobule 1 L | 1.97 | 0.08 | 1.88 | 0.10 | 19.1 | 0.000 | 0.000 | 0.189 | 1.00 | 0.319 | 0.763 | 0.012 |
| Dorsal Attention B: Post Central 1 L | 1.89 | 0.09 | 1.80 | 0.11 | 12.1 | 0.001 | 0.001 | 0.129 | 2.71 | 0.103 | 0.619 | 0.032 |
| Dorsal Attention B: Post Central 2 L | 1.87 | 0.18 | 1.72 | 0.17 | 14.2 | 0.000 | 0.001 | 0.148 | 0.02 | 0.882 | 0.989 | 0.000 |
| Dorsal Attention B: Post Central 3 L | 1.87 | 0.12 | 1.78 | 0.12 | 9.6 | 0.003 | 0.004 | 0.105 | 0.01 | 0.943 | 0.989 | 0.000 |
| Dorsal Attention B: Frontal Eye Fields 1 L | 1.87 | 0.08 | 1.80 | 0.11 | 8.1 | 0.006 | 0.008 | 0.090 | 1.75 | 0.190 | 0.682 | 0.021 |
| Salience Ventral Attention A: Parietal Operculum | 2.08 | 0.06 | 1.98 | 0.07 | 37.6 | 0.000 | 0.000 | 0.314 | 0.92 | 0.341 | 0.793 | 0.011 |
| Salience Ventral Attention A: Insula: 1 L | 2.18 | 0.06 | 2.03 | 0.10 | 68.4 | 0.000 | 0.000 | 0.455 | 1.48 | 0.228 | 0.682 | 0.018 |
| Salience Ventral Attention A: Insula: 2 L | 2.36 | 0.06 | 2.27 | 0.07 | 29.7 | 0.000 | 0.000 | 0.266 | 0.81 | 0.370 | 0.796 | 0.010 |
| Salience Ventral Attention A: Parietal Medial 1 L | 2.02 | 0.06 | 1.95 | 0.11 | 11.7 | 0.001 | 0.002 | 0.124 | 2.65 | 0.108 | 0.619 | 0.031 |
| Salience Ventral Attention A: Frontal Medial 1 L | 2.09 | 0.09 | 1.99 | 0.14 | 12.8 | 0.001 | 0.001 | 0.135 | 1.80 | 0.183 | 0.682 | 0.021 |
| Salience Ventral Attention B: Lateral Prefrontal Co | 2.02 | 0.09 | 1.95 | 0.09 | 11.1 | 0.001 | 0.002 | 0.119 | 4.07 | 0.047 | 0.586 | 0.047 |
| Salience Ventral Attention B: Medial Posterior Pre | 2.00 | 0.08 | 1.91 | 0.11 | 14.7 | 0.000 | 0.000 | 0.152 | 0.00 | 0.974 | 0.990 | 0.000 |
| Limbic A Temporal Pole 1 L | 2.18 | 0.05 | 2.13 | 0.07 | 12.6 | 0.001 | 0.001 | 0.133 | 0.03 | 0.869 | 0.989 | 0.000 |
| Limbic A: Temporal Pole 2 L | 2.23 | 0.06 | 2.21 | 0.07 | 0.7 | 0.417 | 0.438 | 0.008 | 1.53 | 0.219 | 0.682 | 0.018 |
| Limbic B: Orbital Frontal Cortex 1 L | 2.22 | 0.05 | 2.23 | 0.06 | 0.0 | 0.967 | 0.967 | 0.000 | 0.36 | 0.550 | 0.901 | 0.004 |
| Control A: Intraparietal Sulcus 1 L | 1.95 | 0.09 | 1.83 | 0.13 | 22.2 | 0.000 | 0.000 | 0.213 | 0.59 | 0.443 | 0.821 | 0.007 |
| Control A: Lateral Prefrontal Cortex 1 L | 1.92 | 0.07 | 1.84 | 0.10 | 18.8 | 0.000 | 0.000 | 0.187 | 0.15 | 0.701 | 0.952 | 0.002 |
| Control A: Lateral Prefrontal Cortex 2 L | 1.95 | 0.10 | 1.89 | 0.10 | 6.2 | 0.015 | 0.019 | 0.070 | 0.30 | 0.585 | 0.929 | 0.004 |
| Control B: Lateral Prefrontal Cortexv 1 L | 2.07 | 0.07 | 2.05 | 0.08 | 1.6 | 0.205 | 0.223 | 0.020 | 0.00 | 0.945 | 0.989 | 0.000 |
| Control C: Precuneus 1 L | 2.17 | 0.09 | 2.04 | 0.12 | 27.8 | 0.000 | 0.000 | 0.253 | 1.85 | 0.178 | 0.682 | 0.022 |
| Control C: Precuneus 2 L | 2.11 | 0.09 | 2.01 | 0.13 | 18.8 | 0.000 | 0.000 | 0.187 | 0.14 | 0.714 | 0.952 | 0.002 |
| Control C: Cingulate Posterior 1 L | 1.93 | 0.11 | 1.93 | 0.14 | 0.0 | 0.824 | 0.833 | 0.001 | 2.87 | 0.094 | 0.619 | 0.034 |
| Default A: Dorsal Prefrontal Cortex 1 L | 2.08 | 0.08 | 1.94 | 0.10 | 44.5 | 0.000 | 0.000 | 0.352 | 1.52 | 0.221 | 0.682 | 0.018 |
| Default A: Precuneus Posterior Cingulate Cortex | 2.10 | 0.06 | 2.06 | 0.08 | 6.9 | 0.010 | 0.014 | 0.078 | 0.24 | 0.625 | 0.947 | 0.003 |
| Default A: Medial Prefrontal Cortex 1 L | 2.12 | 0.06 | 2.07 | 0.06 | 12.3 | 0.001 | 0.001 | 0.130 | 4.74 | 0.032 | 0.469 | 0.055 |
| Default B: Temp 1 L | 2.17 | 0.06 | 2.14 | 0.06 | 6.0 | 0.017 | 0.021 | 0.068 | 1.52 | 0.221 | 0.682 | 0.018 |
| Default B: Temp 2 L | 2.15 | 0.08 | 2.07 | 0.08 | 26.4 | 0.000 | 0.000 | 0.243 | 0.41 | 0.523 | 0.890 | 0.005 |
| Default B: Inferior Parietal Lobule 1 L | 1.98 | 0.06 | 1.91 | 0.08 | 18.5 | 0.000 | 0.000 | 0.184 | 0.00 | 0.980 | 0.990 | 0.000 |
| Default B: Dorsal Prefrontal Cortex 1 L | 2.11 | 0.08 | 1.98 | 0.09 | 44.1 | 0.000 | 0.000 | 0.350 | 4.71 | 0.033 | 0.469 | 0.054 |
| Default B: Lateral Prefrontal Cortex 1 L | 1.97 | 0.11 | 1.91 | 0.11 | 5.2 | 0.025 | 0.030 | 0.060 | 1.00 | 0.320 | 0.763 | 0.012 |
| Default B: Ventral Prefrontal Cortex 1 L | 2.22 | 0.06 | 2.11 | 0.10 | 38.0 | 0.000 | 0.000 | 0.317 | 0.21 | 0.645 | 0.947 | 0.003 |
| Default B: Ventral Prefrontal Cortex 2 L | 2.13 | 0.05 | 2.05 | 0.08 | 29.6 | 0.000 | 0.000 | 0.265 | 0.34 | 0.561 | 0.905 | 0.004 |
| Default C: RetroSuperior Parietal Lobuleenial 1 L | 2.28 | 0.06 | 2.20 | 0.12 | 11.9 | 0.001 | 0.001 | 0.127 | 0.09 | 0.763 | 0.978 | 0.001 |
| Default C: Parahippocampal Cortex 1 L | 2.35 | 0.09 | 2.25 | 0.13 | 19.4 | 0.000 | 0.000 | 0.191 | 1.94 | 0.168 | 0.682 | 0.023 |
| Temporal Parietal 1 L | 2.19 | 0.09 | 2.11 | 0.08 | 15.1 | 0.000 | 0.000 | 0.156 | 0.19 | 0.663 | 0.947 | 0.002 |
| Visual Central: Extra Striate Cortex 1 R | 2.16 | 0.08 | 2.16 | 0.11 | 0.1 | 0.808 | 0.824 | 0.001 | 0.37 | 0.547 | 0.901 | 0.004 |
| Visual Central: Extra Striate Cortex 2 R | 2.05 | 0.05 | 1.96 | 0.07 | 46.4 | 0.000 | 0.000 | 0.361 | 0.66 | 0.421 | 0.809 | 0.008 |
| Visual Central: Extra Striate Cortex 3 R | 1.96 | 0.09 | 1.89 | 0.09 | 11.9 | 0.001 | 0.001 | 0.127 | 0.04 | 0.843 | 0.989 | 0.000 |
| Visual Peripheral: Striate Cortex Calcarine 1 R | 2.04 | 0.08 | 1.99 | 0.07 | 8.3 | 0.005 | 0.007 | 0.092 | 0.59 | 0.443 | 0.821 | 0.007 |
| Visual Peripheral: Extra Striate Inferior 1 R | 2.06 | 0.09 | 1.99 | 0.08 | 14.7 | 0.000 | 0.000 | 0.152 | 0.01 | 0.935 | 0.989 | 0.000 |
| Visual Peripheral: Extra Striate Superior 1 R | 1.95 | 0.08 | 1.85 | 0.08 | 29.8 | 0.000 | 0.000 | 0.267 | 0.17 | 0.683 | 0.952 | 0.002 |
| Somatomotor A: 1 R | 2.17 | 0.11 | 2.04 | 0.15 | 15.0 | 0.000 | 0.000 | 0.154 | 5.31 | 0.024 | 0.469 | 0.061 |
| Somatomotor A: 2 R | 2.04 | 0.08 | 1.95 | 0.14 | 9.8 | 0.002 | 0.004 | 0.107 | 4.73 | 0.033 | 0.469 | 0.055 |
| Somatomotor A: 3 R | 1.83 | 0.14 | 1.73 | 0.14 | 8.6 | 0.004 | 0.006 | 0.095 | 2.71 | 0.104 | 0.619 | 0.032 |
| Somatomotor A: 4 R | 2.02 | 0.08 | 1.91 | 0.11 | 21.0 | 0.000 | 0.000 | 0.204 | 5.68 | 0.019 | 0.469 | 0.065 |
| Somatomotor B: Auditory 1 R | 1.99 | 0.06 | 1.90 | 0.09 | 27.8 | 0.000 | 0.000 | 0.253 | 0.23 | 0.635 | 0.947 | 0.003 |
| Somatomotor B: S2 1 R | 2.25 | 0.07 | 2.10 | 0.11 | 51.0 | 0.000 | 0.000 | 0.384 | 2.50 | 0.118 | 0.619 | 0.030 |
| Somatomotor B: S2 2 R | 2.17 | 0.07 | 1.93 | 0.14 | 83.3 | 0.000 | 0.000 | 0.504 | 1.70 | 0.196 | 0.682 | 0.020 |
| Somatomotor B: Central 1 R | 1.99 | 0.08 | 1.92 | 0.10 | 8.4 | 0.005 | 0.007 | 0.093 | 1.36 | 0.247 | 0.694 | 0.016 |
| Dorsal Attention A: Temporal Occipital 1 R | 2.13 | 0.07 | 2.09 | 0.08 | 5.0 | 0.029 | 0.034 | 0.057 | 0.02 | 0.881 | 0.989 | 0.000 |
| Dorsal Attention A: Parietal Occipital 1 R | 2.08 | 0.07 | 2.02 | 0.08 | 12.1 | 0.001 | 0.001 | 0.128 | 1.30 | 0.257 | 0.694 | 0.016 |
| Dorsal Attention A: Superior Parietal Lobule 1 R | 1.81 | 0.09 | 1.70 | 0.11 | 16.9 | 0.000 | 0.000 | 0.171 | 1.31 | 0.256 | 0.694 | 0.016 |
| Dorsal Attention B: Post Central 1 R | 2.05 | 0.13 | 1.93 | 0.10 | 19.4 | 0.000 | 0.000 | 0.191 | 0.20 | 0.655 | 0.947 | 0.002 |
| Dorsal Attention B: Post Central 2 R | 1.98 | 0.11 | 1.89 | 0.14 | 8.1 | 0.006 | 0.008 | 0.090 | 1.86 | 0.176 | 0.682 | 0.022 |
| Dorsal Attention B: Frontal Eye Fields 1 R | 1.82 | 0.09 | 1.72 | 0.11 | 17.7 | 0.000 | 0.000 | 0.177 | 3.55 | 0.063 | 0.619 | 0.041 |
| Salience Ventral Attention A: Parietal Operculum | 2.17 | 0.06 | 2.03 | 0.08 | 71.1 | 0.000 | 0.000 | 0.464 | 0.06 | 0.812 | 0.989 | 0.001 |
| Salience Ventral Attention A: Insula: 1 R | 2.26 | 0.06 | 2.21 | 0.06 | 12.2 | 0.001 | 0.001 | 0.130 | 0.80 | 0.374 | 0.796 | 0.010 |
| Salience Ventral Attention A: Parietal Medial 1 R | 2.07 | 0.07 | 2.02 | 0.11 | 5.2 | 0.025 | 0.030 | 0.060 | 1.59 | 0.211 | 0.682 | 0.019 |
| Salience Ventral Attention A: Frontal Medial 1 R | 2.09 | 0.09 | 1.99 | 0.13 | 12.2 | 0.001 | 0.001 | 0.130 | 2.54 | 0.115 | 0.619 | 0.030 |
| Salience Ventral Attention B: Inferior Parietal Lobu | 1.91 | 0.08 | 1.82 | 0.11 | 15.5 | 0.000 | 0.000 | 0.159 | 0.02 | 0.886 | 0.989 | 0.000 |
| Salience Ventral Attention B: Lateral Prefrontal Co | 2.02 | 0.08 | 1.97 | 0.09 | 6.7 | 0.011 | 0.014 | 0.076 | 0.03 | 0.863 | 0.989 | 0.000 |
| Salience Ventral Attention B: Medial Posterior Pre | 2.04 | 0.09 | 1.91 | 0.13 | 21.2 | 0.000 | 0.000 | 0.206 | 7.33 | 0.008 | 0.469 | 0.082 |
| Limbic B: Orbital Frontal Cortex 0 R | 2.24 | 0.04 | 2.20 | 0.06 | 6.8 | 0.011 | 0.014 | 0.076 | 0.00 | 0.990 | 0.990 | 0.000 |
| Limbic A: Temporal Pole 1 R | 2.24 | 0.05 | 2.21 | 0.07 | 2.0 | 0.161 | 0.177 | 0.024 | 0.44 | 0.509 | 0.890 | 0.005 |
| Control A: Intraparietal Sulcus 1 R | 1.72 | 0.15 | 1.67 | 0.14 | 4.2 | 0.044 | 0.051 | 0.049 | 3.00 | 0.087 | 0.619 | 0.035 |
| Control A: Lateral Prefrontal Cortex 1 R | 1.91 | 0.09 | 1.83 | 0.11 | 13.0 | 0.001 | 0.001 | 0.137 | 0.00 | 0.986 | 0.990 | 0.000 |
| Control A: Lateral Prefrontal Cortex 2 R | 1.83 | 0.08 | 1.77 | 0.10 | 9.3 | 0.003 | 0.005 | 0.101 | 0.08 | 0.782 | 0.978 | 0.001 |
| Control B: Temporal 1 R | 2.21 | 0.06 | 2.18 | 0.07 | 6.7 | 0.011 | 0.015 | 0.075 | 2.97 | 0.089 | 0.619 | 0.035 |
| Control B: inferior parietal lobule 1 R | 2.02 | 0.08 | 1.90 | 0.10 | 35.2 | 0.000 | 0.000 | 0.300 | 0.12 | 0.729 | 0.959 | 0.001 |
| Control B: Lateral Prefrontal Cortexd 1 R | 2.05 | 0.09 | 1.94 | 0.10 | 21.2 | 0.000 | 0.000 | 0.206 | 1.45 | 0.232 | 0.682 | 0.017 |
| Control B: Lateral Prefrontal Cortexv 1 R | 2.01 | 0.05 | 1.99 | 0.07 | 2.7 | 0.106 | 0.119 | 0.032 | 0.14 | 0.710 | 0.952 | 0.002 |
| Control C: Cingulate Posterior 1 R | 2.03 | 0.06 | 2.01 | 0.11 | 0.8 | 0.383 | 0.408 | 0.009 | 0.70 | 0.404 | 0.809 | 0.008 |
| Control C: Precuneus 1 R | 2.17 | 0.06 | 2.06 | 0.11 | 26.5 | 0.000 | 0.000 | 0.244 | 2.91 | 0.092 | 0.619 | 0.034 |
| Default A: Inferior Parietal Lobule 1 R | 1.99 | 0.10 | 1.91 | 0.10 | 14.9 | 0.000 | 0.000 | 0.154 | 0.80 | 0.373 | 0.796 | 0.010 |
| Default A: Dorsal Prefrontal Cortex 1 R | 1.94 | 0.10 | 1.83 | 0.09 | 27.7 | 0.000 | 0.000 | 0.252 | 1.56 | 0.215 | 0.682 | 0.019 |
| Default A: Precuneus Posterior Cingulate Cortex | 2.17 | 0.06 | 2.16 | 0.07 | 0.1 | 0.801 | 0.824 | 0.001 | 0.16 | 0.690 | 0.952 | 0.002 |
| Default A: Medial Prefrontal Cortex 1 R | 2.20 | 0.05 | 2.12 | 0.07 | 25.4 | 0.000 | 0.000 | 0.236 | 3.35 | 0.071 | 0.619 | 0.039 |
| Default B: Dorsal Prefrontal Cortex 1 R | 2.07 | 0.08 | 1.97 | 0.08 | 25.3 | 0.000 | 0.000 | 0.235 | 2.02 | 0.159 | 0.682 | 0.024 |
| Default B: Ventral Prefrontal Cortex 1 R | 2.18 | 0.06 | 2.11 | 0.07 | 24.4 | 0.000 | 0.000 | 0.229 | 0.76 | 0.386 | 0.804 | 0.009 |
| Default B: Ventral Prefrontal Cortex 2 R | 2.16 | 0.06 | 2.04 | 0.09 | 52.6 | 0.000 | 0.000 | 0.391 | 0.68 | 0.413 | 0.809 | 0.008 |
| Default C: RetroSuperior Parietal Lobuleenial 1 R | 2.36 | 0.07 | 2.28 | 0.09 | 19.2 | 0.000 | 0.000 | 0.190 | 0.04 | 0.841 | 0.989 | 0.000 |
| Default C: Parahippocampal Cortex 1 R | 2.31 | 0.07 | 2.29 | 0.08 | 2.1 | 0.153 | 0.170 | 0.025 | 1.22 | 0.272 | 0.712 | 0.015 |
| Temporal Parietal 1 R | 2.10 | 0.08 | 2.01 | 0.07 | 26.5 | 0.000 | 0.000 | 0.244 | 0.08 | 0.780 | 0.978 | 0.001 |
| Temporal Parietal 2 R | 2.09 | 0.06 | 2.01 | 0.07 | 31.8 | 0.000 | 0.000 | 0.279 | 0.41 | 0.525 | 0.890 | 0.005 |
| Temporal Parietal 3 R | 1.95 | 0.10 | 1.91 | 0.08 | 4.6 | 0.034 | 0.040 | 0.053 | 0.11 | 0.740 | 0.961 | 0.001 |

**Table S8. Whole sample and younger and older adult group regional fPET standard deviation.** Mean and standard deviation of 100 region $fPET_{SD}$ values and GLMs of age group differences.

| | Younger | | Older | | Younger vs older | | | | Head motion | | | |
|---|---|---|---|---|---|---|---|---|---|---|---|---|
| | Mean | SD | Mean | SD | F | p | p-FDR | η2p | F | p | p-FDR | η2p |
| Visual Central: Extra Striate Cortex 1 L | 1852.5 | 155.7 | 1776.4 | 237.5 | 1.8 | 0.182 | 0.196 | 0.022 | 2.54 | 0.115 | 0.115 | 0.030 |
| Visual Central: Extra Striate Cortex 2 L | 1683.8 | 164.9 | 1601.4 | 210.6 | 2.0 | 0.161 | 0.175 | 0.024 | 7.47 | 0.008 | 0.014 | 0.084 |
| Visual Central: Striate Cortex 1 L | 1975.2 | 190.5 | 1869.7 | 254.0 | 2.7 | 0.102 | 0.114 | 0.032 | 4.86 | 0.030 | 0.032 | 0.056 |
| Visual Central: Extra Striate Cortex 3 L | 1648.0 | 149.6 | 1558.8 | 173.3 | 3.6 | 0.060 | 0.073 | 0.042 | 9.57 | 0.003 | 0.013 | 0.105 |
| Visual Peripheral: Extra Striate Inferior 1 L | 1945.4 | 164.9 | 1828.6 | 208.4 | 5.2 | 0.025 | 0.037 | 0.060 | 7.07 | 0.009 | 0.015 | 0.079 |
| Visual Peripheral: Striate Cortex Calcarine 1 L | 1948.7 | 166.1 | 1822.2 | 206.9 | 6.1 | 0.016 | 0.027 | 0.069 | 10.04 | 0.002 | 0.013 | 0.109 |
| Visual Peripheral: Extra Striate CortexSup 1 L | 1817.7 | 193.1 | 1710.9 | 206.2 | 3.3 | 0.074 | 0.087 | 0.038 | 10.98 | 0.001 | 0.013 | 0.118 |
| Somatomotor A: 1 L | 1970.6 | 246.9 | 1814.4 | 322.2 | 4.0 | 0.050 | 0.063 | 0.046 | 4.96 | 0.029 | 0.031 | 0.057 |
| Somatomotor A: 2 L | 1995.4 | 246.9 | 1815.1 | 307.3 | 6.1 | 0.016 | 0.027 | 0.069 | 5.02 | 0.028 | 0.030 | 0.058 |
| Somatomotor B: Auditory 1 L | 1759.5 | 174.0 | 1683.6 | 194.2 | 1.7 | 0.191 | 0.203 | 0.021 | 7.27 | 0.008 | 0.015 | 0.081 |
| Somatomotor B: S2 1 L | 2079.3 | 203.2 | 1893.0 | 263.8 | 8.9 | 0.004 | 0.010 | 0.098 | 12.24 | 0.001 | 0.013 | 0.130 |
| Somatomotor B: S2 2 L | 1602.2 | 147.0 | 1461.6 | 183.1 | 10.7 | 0.002 | 0.007 | 0.115 | 11.08 | 0.001 | 0.013 | 0.119 |
| Somatomotor B: Central 1 L | 1829.7 | 189.7 | 1715.8 | 245.4 | 3.2 | 0.077 | 0.088 | 0.038 | 7.85 | 0.006 | 0.014 | 0.087 |
| Dorsal Attention A: Temporal Occipital 1 L | 1560.0 | 115.5 | 1452.5 | 156.3 | 9.2 | 0.003 | 0.009 | 0.101 | 6.57 | 0.012 | 0.018 | 0.074 |
| Dorsal Attention A: Parietal Occipital 1 L | 1574.9 | 138.9 | 1484.8 | 161.3 | 4.4 | 0.038 | 0.052 | 0.051 | 10.82 | 0.001 | 0.013 | 0.117 |
| Dorsal Attention A: Superior Parietal Lobule 1 L | 1777.6 | 184.4 | 1633.0 | 245.0 | 6.1 | 0.016 | 0.027 | 0.069 | 7.94 | 0.006 | 0.014 | 0.088 |
| Dorsal Attention B: Post Central 1 L | 1678.6 | 177.1 | 1518.6 | 220.3 | 9.7 | 0.003 | 0.009 | 0.106 | 7.09 | 0.009 | 0.015 | 0.080 |
| Dorsal Attention B: Post Central 2 L | 1895.5 | 212.5 | 1723.0 | 269.9 | 7.4 | 0.008 | 0.018 | 0.082 | 6.26 | 0.014 | 0.020 | 0.071 |
| Dorsal Attention B: Post Central 3 L | 1799.0 | 218.7 | 1598.8 | 290.6 | 9.2 | 0.003 | 0.009 | 0.101 | 5.74 | 0.019 | 0.023 | 0.065 |
| Dorsal Attention B: Frontal Eye Fields 1 L | 1968.9 | 238.9 | 1780.3 | 303.8 | 7.1 | 0.009 | 0.019 | 0.079 | 5.03 | 0.028 | 0.030 | 0.058 |
| Salience Ventral Attention A: Parietal Operculum 1 | 1646.7 | 181.3 | 1495.2 | 207.8 | 9.0 | 0.004 | 0.010 | 0.099 | 7.99 | 0.006 | 0.014 | 0.089 |
| Salience Ventral Attention A: Insula: 1 L | 1579.8 | 137.3 | 1469.1 | 170.2 | 7.2 | 0.009 | 0.018 | 0.081 | 8.82 | 0.004 | 0.013 | 0.097 |
| Salience Ventral Attention A: Insula: 2 L | 1918.8 | 191.2 | 1701.4 | 217.2 | 18.8 | 0.000 | 0.001 | 0.186 | 7.36 | 0.008 | 0.015 | 0.082 |
| Salience Ventral Attention A: Parietal Medial 1 L | 2080.3 | 243.8 | 1845.5 | 294.4 | 11.5 | 0.001 | 0.006 | 0.123 | 9.26 | 0.003 | 0.013 | 0.102 |
| Salience Ventral Attention A: Frontal Medial 1 L | 1897.4 | 232.1 | 1712.7 | 307.4 | 6.6 | 0.012 | 0.023 | 0.075 | 5.88 | 0.018 | 0.022 | 0.067 |
| Salience Ventral Attention B: Lateral Prefrontal Cor | 1788.4 | 191.9 | 1599.1 | 234.9 | 12.4 | 0.001 | 0.004 | 0.132 | 5.54 | 0.021 | 0.025 | 0.063 |
| Salience Ventral Attention B: Medial Posterior Prefr | 1869.6 | 188.9 | 1666.6 | 270.1 | 11.5 | 0.001 | 0.006 | 0.123 | 8.70 | 0.004 | 0.013 | 0.096 |
| Limbic A Temporal Pole 1 L | 1758.8 | 162.8 | 1548.3 | 179.1 | 26.2 | 0.000 | 0.000 | 0.242 | 7.07 | 0.009 | 0.015 | 0.079 |
| Limbic A: Temporal Pole 2 L | 1310.2 | 105.5 | 1228.0 | 129.9 | 7.1 | 0.009 | 0.019 | 0.080 | 5.31 | 0.024 | 0.027 | 0.061 |
| Limbic B: Orbital Frontal Cortex 1 L | 1577.2 | 128.6 | 1482.0 | 161.0 | 6.1 | 0.016 | 0.027 | 0.069 | 5.85 | 0.018 | 0.022 | 0.067 |
| Control A: Intraparietal Sulcus 1 L | 1824.0 | 200.4 | 1641.2 | 252.7 | 9.7 | 0.003 | 0.009 | 0.106 | 7.23 | 0.009 | 0.015 | 0.081 |
| Control A: Lateral Prefrontal Cortex 1 L | 1797.6 | 200.1 | 1624.5 | 239.6 | 9.2 | 0.003 | 0.009 | 0.101 | 7.26 | 0.009 | 0.015 | 0.081 |
| Control A: Lateral Prefrontal Cortex 2 L | 1850.3 | 193.5 | 1663.9 | 247.8 | 10.7 | 0.002 | 0.007 | 0.116 | 7.65 | 0.007 | 0.014 | 0.085 |
| Control B: Lateral Prefrontal Cortexv 1 L | 1657.9 | 152.9 | 1475.7 | 187.5 | 18.8 | 0.000 | 0.001 | 0.186 | 7.64 | 0.007 | 0.014 | 0.085 |
| Control C: Precuneus 1 L | 1881.4 | 197.8 | 1780.0 | 240.8 | 2.5 | 0.118 | 0.130 | 0.030 | 5.90 | 0.017 | 0.022 | 0.067 |
| Control C: Precuneus 2 L | 1759.3 | 212.2 | 1610.6 | 263.8 | 5.2 | 0.026 | 0.037 | 0.059 | 7.69 | 0.007 | 0.014 | 0.086 |
| Control C: Cingulate Posterior 1 L | 1994.1 | 248.2 | 1742.2 | 284.9 | 13.9 | 0.000 | 0.003 | 0.145 | 12.33 | 0.001 | 0.013 | 0.131 |
| Default A: Dorsal Prefrontal Cortex 1 L | 1960.2 | 236.8 | 1748.2 | 309.7 | 8.7 | 0.004 | 0.010 | 0.095 | 8.14 | 0.005 | 0.014 | 0.090 |
| Default A: Precuneus Posterior Cingulate Cortex1 | 1920.5 | 225.4 | 1743.4 | 243.0 | 8.4 | 0.005 | 0.011 | 0.093 | 8.38 | 0.005 | 0.013 | 0.093 |
| Default A: Medial Prefrontal Cortex 1 L | 1644.7 | 165.1 | 1463.5 | 189.6 | 16.9 | 0.000 | 0.001 | 0.171 | 8.53 | 0.005 | 0.013 | 0.094 |
| Default B: Temp 1 L | 1451.1 | 122.6 | 1332.5 | 138.0 | 13.1 | 0.001 | 0.004 | 0.138 | 7.34 | 0.008 | 0.015 | 0.082 |
| Default B: Temp 2 L | 1653.8 | 142.9 | 1555.1 | 166.2 | 5.6 | 0.020 | 0.031 | 0.064 | 6.78 | 0.011 | 0.016 | 0.076 |
| Default B: Inferior Parietal Lobule 1 L | 1593.4 | 172.1 | 1451.6 | 200.6 | 8.4 | 0.005 | 0.011 | 0.093 | 8.55 | 0.004 | 0.013 | 0.094 |
| Default B: Dorsal Prefrontal Cortex 1 L | 1704.9 | 194.5 | 1519.1 | 248.5 | 10.8 | 0.002 | 0.007 | 0.116 | 6.16 | 0.015 | 0.020 | 0.070 |
| Default B: Lateral Prefrontal Cortex 1 L | 1830.8 | 232.4 | 1670.6 | 290.7 | 5.1 | 0.027 | 0.038 | 0.059 | 5.92 | 0.017 | 0.022 | 0.067 |
| Default B: Ventral Prefrontal Cortex 1 L | 1666.3 | 168.9 | 1472.2 | 166.6 | 22.8 | 0.000 | 0.000 | 0.217 | 10.70 | 0.002 | 0.013 | 0.115 |
| Default B: Ventral Prefrontal Cortex 2 L | 1632.4 | 163.2 | 1470.4 | 181.3 | 14.1 | 0.000 | 0.003 | 0.147 | 7.91 | 0.006 | 0.014 | 0.088 |
| Default C: RetroSuperior Parietal Lobuleenial 1 L | 1917.0 | 197.8 | 1768.3 | 220.4 | 7.0 | 0.010 | 0.019 | 0.079 | 9.84 | 0.002 | 0.013 | 0.107 |
| Default C: Parahippocampal Cortex 1 L | 1635.3 | 133.9 | 1523.7 | 157.5 | 8.8 | 0.004 | 0.010 | 0.097 | 6.09 | 0.016 | 0.021 | 0.069 |
| Temporal Parietal 1 L | 1575.8 | 128.5 | 1457.2 | 159.7 | 10.0 | 0.002 | 0.008 | 0.109 | 9.11 | 0.003 | 0.013 | 0.100 |
| Visual Central: Extra Striate Cortex 1 R | 1854.2 | 152.9 | 1784.5 | 215.2 | 1.5 | 0.228 | 0.238 | 0.018 | 4.85 | 0.030 | 0.032 | 0.056 |
| Visual Central: Extra Striate Cortex 2 R | 1768.2 | 175.9 | 1701.8 | 232.6 | 1.0 | 0.332 | 0.335 | 0.011 | 5.03 | 0.028 | 0.030 | 0.058 |
| Visual Central: Extra Striate Cortex 3 R | 1666.6 | 148.4 | 1596.4 | 191.8 | 1.4 | 0.234 | 0.241 | 0.017 | 11.12 | 0.001 | 0.013 | 0.119 |
| Visual Peripheral: Striate Cortex Calcarine 1 R | 1983.5 | 178.5 | 1870.8 | 241.2 | 3.8 | 0.055 | 0.068 | 0.044 | 4.53 | 0.036 | 0.037 | 0.052 |
| Visual Peripheral: Extra Striate Inferior 1 R | 1882.5 | 160.4 | 1748.3 | 201.2 | 7.4 | 0.008 | 0.017 | 0.083 | 13.04 | 0.001 | 0.013 | 0.137 |
| Visual Peripheral: Extra Striate Superior 1 R | 1721.3 | 176.2 | 1618.2 | 193.0 | 4.2 | 0.044 | 0.056 | 0.049 | 5.47 | 0.022 | 0.025 | 0.063 |
| Somatomotor A: 1 R | 2037.5 | 264.0 | 1883.8 | 312.6 | 3.7 | 0.057 | 0.070 | 0.043 | 5.22 | 0.025 | 0.028 | 0.060 |
| Somatomotor A: 2 R | 1977.8 | 235.0 | 1819.0 | 305.9 | 4.5 | 0.037 | 0.051 | 0.052 | 6.30 | 0.014 | 0.020 | 0.071 |
| Somatomotor A: 3 R | 1961.3 | 248.1 | 1815.4 | 301.3 | 3.6 | 0.061 | 0.073 | 0.042 | 5.86 | 0.018 | 0.022 | 0.067 |
| Somatomotor A: 4 R | 1985.8 | 244.8 | 1815.3 | 318.7 | 5.0 | 0.028 | 0.040 | 0.057 | 5.49 | 0.022 | 0.025 | 0.063 |
| Somatomotor B: Auditory 1 R | 1748.7 | 149.0 | 1695.7 | 204.0 | 0.5 | 0.480 | 0.480 | 0.006 | 9.43 | 0.003 | 0.013 | 0.103 |
| Somatomotor B: S2 1 R | 1924.3 | 177.6 | 1777.1 | 245.2 | 6.3 | 0.014 | 0.025 | 0.072 | 10.41 | 0.002 | 0.013 | 0.113 |
| Somatomotor B: S2 2 R | 1550.4 | 141.9 | 1459.4 | 204.6 | 3.2 | 0.079 | 0.090 | 0.037 | 7.73 | 0.007 | 0.014 | 0.086 |
| Somatomotor B: Central 1 R | 1792.4 | 190.1 | 1710.4 | 269.7 | 1.0 | 0.309 | 0.316 | 0.013 | 7.45 | 0.008 | 0.014 | 0.083 |
| Dorsal Attention A: Temporal Occipital 1 R | 1544.8 | 121.6 | 1468.1 | 159.1 | 3.9 | 0.052 | 0.065 | 0.045 | 5.09 | 0.027 | 0.030 | 0.058 |
| Dorsal Attention A: Parietal Occipital 1 R | 1557.7 | 131.4 | 1490.9 | 173.6 | 1.7 | 0.193 | 0.203 | 0.021 | 11.23 | 0.001 | 0.013 | 0.120 |
| Dorsal Attention A: Superior Parietal Lobule 1 R | 1759.3 | 192.3 | 1634.9 | 237.8 | 4.3 | 0.042 | 0.055 | 0.049 | 7.50 | 0.008 | 0.014 | 0.084 |
| Dorsal Attention B: Post Central 1 R | 1913.1 | 213.7 | 1761.9 | 258.8 | 5.5 | 0.022 | 0.032 | 0.063 | 7.80 | 0.007 | 0.014 | 0.087 |
| Dorsal Attention B: Post Central 2 R | 1767.9 | 216.3 | 1601.6 | 285.1 | 6.1 | 0.016 | 0.027 | 0.069 | 6.35 | 0.014 | 0.019 | 0.072 |
| Dorsal Attention B: Frontal Eye Fields 1 R | 1985.6 | 248.9 | 1786.7 | 323.2 | 6.6 | 0.012 | 0.023 | 0.074 | 8.49 | 0.005 | 0.013 | 0.094 |
| Salience Ventral Attention A: Parietal Operculum 1 | 1630.5 | 154.5 | 1481.8 | 214.7 | 9.3 | 0.003 | 0.009 | 0.101 | 8.90 | 0.004 | 0.013 | 0.098 |
| Salience Ventral Attention A: Insula: 1 R | 1743.4 | 156.5 | 1590.9 | 207.3 | 10.2 | 0.002 | 0.008 | 0.111 | 9.71 | 0.003 | 0.013 | 0.106 |
| Salience Ventral Attention A: Parietal Medial 1 R | 2069.5 | 244.8 | 1844.2 | 290.7 | 10.5 | 0.002 | 0.007 | 0.114 | 10.04 | 0.002 | 0.013 | 0.109 |
| Salience Ventral Attention A: Frontal Medial 1 R | 1921.9 | 240.2 | 1735.5 | 305.6 | 6.3 | 0.014 | 0.025 | 0.072 | 8.61 | 0.004 | 0.013 | 0.095 |
| Salience Ventral Attention B: Inferior Parietal Lobule | 1578.4 | 172.0 | 1446.3 | 225.5 | 5.8 | 0.018 | 0.029 | 0.066 | 9.43 | 0.003 | 0.013 | 0.103 |
| Salience Ventral Attention B: Lateral Prefrontal Cor | 1763.2 | 209.8 | 1578.9 | 252.0 | 9.4 | 0.003 | 0.009 | 0.103 | 8.71 | 0.004 | 0.013 | 0.096 |
| Salience Ventral Attention B: Medial Posterior Prefr | 1892.4 | 231.4 | 1638.0 | 269.9 | 16.4 | 0.000 | 0.001 | 0.167 | 12.43 | 0.001 | 0.013 | 0.132 |
| Limbic B: Orbital Frontal Cortex 0 R | 1724.6 | 163.6 | 1509.1 | 185.6 | 26.2 | 0.000 | 0.000 | 0.242 | 7.54 | 0.007 | 0.014 | 0.084 |
| Limbic A: Temporal Pole 1 R | 1380.5 | 111.7 | 1275.8 | 134.4 | 11.4 | 0.001 | 0.006 | 0.122 | 5.52 | 0.021 | 0.025 | 0.063 |
| Control A: Intraparietal Sulcus 1 R | 1859.8 | 224.1 | 1712.9 | 273.5 | 4.5 | 0.037 | 0.051 | 0.052 | 7.71 | 0.007 | 0.014 | 0.086 |
| Control A: Lateral Prefrontal Cortex 1 R | 1759.5 | 194.4 | 1608.6 | 218.7 | 7.3 | 0.008 | 0.018 | 0.082 | 12.73 | 0.001 | 0.013 | 0.134 |
| Control A: Lateral Prefrontal Cortex 2 R | 1883.6 | 205.8 | 1695.3 | 247.5 | 10.5 | 0.002 | 0.007 | 0.113 | 7.15 | 0.009 | 0.015 | 0.080 |
| Control B: Temporal 1 R | 1502.5 | 119.0 | 1397.1 | 155.9 | 9.0 | 0.004 | 0.010 | 0.099 | 4.38 | 0.040 | 0.040 | 0.051 |
| Control B: inferior parietal lobule 1 R | 1617.2 | 186.2 | 1497.2 | 219.8 | 4.3 | 0.041 | 0.055 | 0.050 | 10.03 | 0.002 | 0.013 | 0.109 |
| Control B: Lateral Prefrontal Cortexd 1 R | 1833.7 | 220.0 | 1677.8 | 266.4 | 5.6 | 0.020 | 0.031 | 0.064 | 7.02 | 0.010 | 0.015 | 0.079 |
| Control B: Lateral Prefrontal Cortexv 1 R | 1628.2 | 168.1 | 1457.0 | 193.3 | 14.2 | 0.000 | 0.003 | 0.148 | 8.45 | 0.005 | 0.013 | 0.093 |
| Control C: Cingulate Posterior 1 R | 2016.9 | 252.6 | 1736.7 | 298.3 | 16.7 | 0.000 | 0.001 | 0.169 | 8.40 | 0.005 | 0.013 | 0.093 |
| Control C: Precuneus 1 R | 1825.8 | 200.4 | 1704.9 | 250.4 | 3.5 | 0.064 | 0.075 | 0.041 | 7.16 | 0.009 | 0.015 | 0.080 |
| Default A: Inferior Parietal Lobule 1 R | 1590.4 | 167.5 | 1499.7 | 221.4 | 2.5 | 0.115 | 0.128 | 0.030 | 5.62 | 0.020 | 0.025 | 0.064 |
| Default A: Dorsal Prefrontal Cortex 1 R | 1972.6 | 226.8 | 1761.3 | 303.9 | 9.3 | 0.003 | 0.009 | 0.102 | 7.19 | 0.009 | 0.015 | 0.081 |
| Default A: Precuneus Posterior Cingulate Cortex 1 | 1920.3 | 213.3 | 1739.9 | 258.0 | 8.3 | 0.005 | 0.011 | 0.092 | 10.01 | 0.002 | 0.013 | 0.109 |
| Default A: Medial Prefrontal Cortex 1 R | 1666.1 | 168.4 | 1458.2 | 200.4 | 21.0 | 0.000 | 0.000 | 0.204 | 11.78 | 0.001 | 0.013 | 0.126 |
| Default B: Dorsal Prefrontal Cortex 1 R | 1734.5 | 201.4 | 1511.8 | 237.4 | 16.6 | 0.000 | 0.001 | 0.168 | 9.31 | 0.003 | 0.013 | 0.102 |
| Default B: Ventral Prefrontal Cortex 1 R | 1634.8 | 160.6 | 1482.0 | 182.2 | 12.4 | 0.001 | 0.004 | 0.132 | 7.67 | 0.007 | 0.014 | 0.086 |
| Default B: Ventral Prefrontal Cortex 2 R | 1617.3 | 162.2 | 1467.9 | 198.4 | 10.6 | 0.002 | 0.007 | 0.115 | 5.58 | 0.021 | 0.025 | 0.064 |
| Default C: RetroSuperior Parietal Lobuleenial 1 R | 1898.0 | 194.5 | 1776.5 | 230.5 | 4.2 | 0.042 | 0.055 | 0.049 | 6.73 | 0.011 | 0.017 | 0.076 |
| Default C: Parahippocampal Cortex 1 R | 1616.5 | 131.2 | 1520.8 | 169.3 | 6.0 | 0.017 | 0.027 | 0.068 | 3.54 | 0.064 | 0.064 | 0.041 |
| Temporal Parietal 1 R | 1429.1 | 116.4 | 1328.2 | 142.6 | 8.8 | 0.004 | 0.010 | 0.097 | 8.67 | 0.004 | 0.013 | 0.096 |
| Temporal Parietal 2 R | 1658.3 | 132.4 | 1555.2 | 176.0 | 5.9 | 0.018 | 0.028 | 0.067 | 9.34 | 0.003 | 0.013 | 0.102 |
| Temporal Parietal 3 R | 1635.5 | 137.3 | 1532.4 | 181.5 | 5.7 | 0.019 | 0.030 | 0.065 | 6.35 | 0.014 | 0.019 | 0.072 |

##### *2.7 Sensitivity Analysis*

To evaluate the stability of the macroscale dynamic states and ensure that our primary findings were not artifacts of our choice of a 4-frame sliding window, we conducted a comprehensive sensitivity analysis across alternative temporal scales. Using the same fPET time-series data, we recalculated sliding-window functional connectivity matrices using window lengths of 6 and 8 frames. To evaluate the clustering stability across these temporal configurations, we executed a comprehensive data validation sweep, generating cluster profiles across an expansion range of k = 2 to k = 8. For each window size and cluster value configuration, the spatial partition quality was quantified using the Within-Cluster Sum of Squares (WCSS; Elbow Method) and the mean Silhouette Coefficient profile. Across all evaluated temporal window scales, cluster validation metrics converged on a 4-state topological architecture as the optimal partition model for the metabolic landscape (Fig S4). WCSS curves displayed similar profiles, while mean Silhouette profiles revealed that expanding the partition past 4 or 5 clusters forced the boundaries into unstructured, negative separation spaces.

The overall chronnectomic architecture remained largely invariant to the chosen window size (Table S9); the sparsely connected configuration (State 4) consistently emerged as the prominent baseline across all scales (54% to 59% global occupancy) and displayed spatial profile stability across window transitions (minimum centroid correlation r = 0.928). Conversely, the highly integrated configuration (State 3) was preserved as a rare, transient state (1.6% to 9.4% occupancy.) A proportional compression of total transitions (62.9 to 45.7 to 32.3) and elongation of dwell times as the window expanded from 4 to 8 frames reflect expected temporal smoothing properties, confirming that the macroscale metabolic dynamics described in our primary analysis are structurally sound, reproducible, and independent of arbitrary window selection.

**Table S9. Stability analysis.** Chronnectomic metric segmentation and centroid spatial stability across sliding-window sensitivity configurations.

| Parameter_Evaluated | 4 Frame Window | 6 Frame Window | 8 Frame Window |
|---|---|---|---|
| Optimal States (k) | k = 4 | k = 4 | k = 4 |
| Mean Transitions (Cohort) | 62.929 | 45.706 | 32.341 |
| State 1 Occupancy (%) | 17.674 | 32.225 | 1.336 |
| State 2 Occupancy (%) | 18.001 | 11.949 | 29.977 |
| State 3 Occupancy (%) | 6.474 | 1.678 | 9.445 |
| State 4 Occupancy (%) | 57.851 | 54.148 | 59.242 |
| State 1 Mean Dwell Time | 1.583 | 2.334 | 4.112 |
| State 2 Mean Dwell Time | 1.610 | 1.994 | 2.822 |
| State 3 Mean Dwell Time | 1.459 | 2.450 | 2.637 |
| State 4 Mean Dwell Time | 3.839 | 5.174 | 9.026 |
| State 1 Centroid Stability (r) | Reference Base | 0.671 | 0.561 |
| State 2 Centroid Stability (r) | Reference Base | 0.708 | 0.725 |
| State 3 Centroid Stability (r) | Reference Base | 0.796 | 0.831 |
| State 4 Centroid Stability (r) | Reference Base | 0.945 | 0.928 |

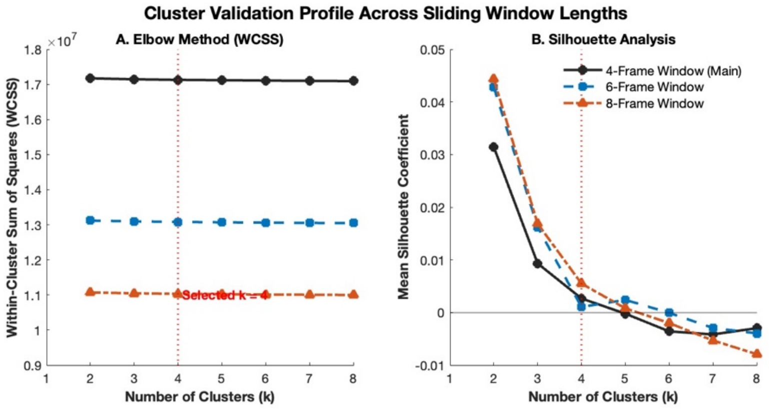


**Figure S4. Cluster validation and optimisation profiles across alternative sliding-window lengths.** Elbow method curves (A) and silhouette coefficient profiles (B) confirming 4-cluster solution.
